# Sustained post-developmental T-bet expression is critical for the maintenance of type one innate lymphoid cells *in vivo*

**DOI:** 10.1101/2021.06.05.447200

**Authors:** J-H Schroeder, LB Roberts, K Meissl, JW Lo, D Hromadová, K Hayes, T Zabinski, E Read, C Moreira Heliodoro, R Reis, JK Howard, RK Grencis, J F Neves, B Strobl, GM Lord

**Affiliations:** Department of Experimental Immunobiology, Division of Transplantation Immunology and Mucosal Biology, King’s College London, London SE1 9RT, UK; Institute of Animal Breeding and Genetics, University of Veterinary Medicine Vienna, Veterinärplatz 1, 1210 Vienna, Austria; Division of Digestive Diseases, Faculty of Medicine, Imperial College London, W12 0NN, UK; School of Biological Sciences, Faculty of Biology, Medicine and Health, Division of Infection, Immunity and Respiratory Medicine, University of Manchester, M13 9PL, UK; Centre for Host-Microbiome Interactions, King’s College London, SE1 9RT, UK; Wellcome Trust Cell Therapies and Regenerative Medicine PhD Programme, London, UK; Department of Diabetes, School of Life Course Sciences, Faculty of Life Sciences and Medicine, King’s College, London, United Kingdom

**Author notes:** These authors contributed equally (LB Roberts, K Meissl, JW Lo). **Corresponding Author:** Graham M Lord MD PhD FMedSci.

**Keywords:** T-bet, Innate lymphoid cells, ILC, intestinal inflammation, mucosal homeostasis

## Abstract

Innate lymphoid cells (ILC) play a significant role in the intestinal immune response and T-bet^+^ CD127^+^ group 1 cells (ILC1) have been linked to the pathogenesis of human inflammatory bowel disease (IBD). However, the functional importance of ILC1 in the context of an intact adaptive immune response has been controversial. In this report we demonstrate that induced depletion of T-bet using a Rosa26-Cre-ERT2 model resulted in the loss of intestinal ILC1, pointing to a post-developmental requirement of T-bet expression for these cells. In contrast, neither colonic lamina propria (cLP) ILC2 nor cLP ILC3 abundance were altered upon induced deletion of T-bet. Mechanistically, we report that STAT1 or STAT4 are not required for intestinal ILC1 development and maintenance. Mice with induced deletion of T-bet and subsequent loss of ILC1 were protected from the induction of severe colitis *in vivo*. Hence, this study provides support for the clinical development of an IBD treatment based on ILC1 depletion via targeting T-bet or its downstream transcriptional targets.

## INTRODUCTION

Innate lymphoid cells (ILC) have been categorised into subsets based on expression of characteristic transcription factors (Bal *et al*., 2020). Group 1 ILC express T box expressed in T cells (T-bet, encoded by *Tbx21*) and include conventional CD127^-^ NK cells (cNK) and CD127^+^ ILC1 (herein referred to as ILC1). CD127^+^ ILC1 do not express the cNK molecules Eomesodermin (Eomes), Granzyme B, Perforin or CD49b. Group 2 ILC (ILC2) express GATA3 while group 3 ILC (ILC3) have a characteristic expression of RORγt. The ILC3 group comprises three subgroups depending on the expression of NKp46 and CCR6. NKp46^+^ ILC3 co-express T-bet and NKp46. CCR6 and NKp46 double-negative (DN) ILC3 have been suggested to be the precursors of these cells (Klose *et al*., 2013). In contrast, lymphoid tissue inducer cells (LTi)-like ILC3 are characterised by expression of CCR6 and do not express T-bet. In addition, CCR6^+^ ILC3 are generated following a developmental pathway that is distinct to other ILC subsets (Ishizuka *et al*., 2016). ILC demonstrate similarities in effector cell functionality with conventional and unconventional T cell subsets; IFNγ and TNF-α are expressed by ILC1, ILC3 produce IL-22, IL-17A and IL-17F, and ILC2 produce IL-13, IL-5, IL-6, IL-9 and IL-4 (Panda *et al*., 2019). It has been reported that ILC2 post-developmental maintenance depends on GATA3, while RORγt inhibition and depletion in mature mice does not alter ILC3 abundance (Yagi *et al*., 2014, Withers *et al*., 2016, Fiancette *et al*., 2021).

We have previously reported that genetic variation at T-bet binding sites is primarily associated with mucosal inflammatory diseases in humans, such as Crohn’s disease and ulcerative colitis (Soderquest *et al*., 2017). Furthermore, IFNγ appears to be the most critical factor driving colitis, while IL-17A and IL-13 play a less important role (Ito *et al*., 2006, Langer *et al*., 2019, Brasseit *et al*., 2018, Karmele *et al*., 2019). Interestingly, these reports are supported by the observation that T-bet^+^ ILC1 are particularly abundant among the total number of ILC in inflamed intestinal lamina propria of Crohn’s disease patients (Bernink *et al*., 2013, Bernink *et al*., 2015 and Fuchs *et al*., 2013, Jowett *et al*., 2020). In mouse models, the generation of ILC1 from an NKp46^+^ ILC3 source has been linked to colitis development (Vonarbourg *et al*., 2010). We have recently observed that NKp46-dependent depletion of T-bet in mice leads to milder dextran sulphate sodium (DSS)-induced colitis, and this may be caused by the depletion of T-bet expressing ILC1 and ILC3 (Garrido Mesa *et al*., 2019). In contrast to studies suggesting that ILC1 promote colitis, we have reported previously that T-bet expression can also protect from colitis, as *Rag2*^-/-^ mice with germline deletion of *Tbx21* can develop spontaneous colitis if *Helicobacter thyphlonius* is present in the gut (Vonarbourg *et al*., 2010, Bernink *et al*., 2013, Bernink *et al*., 2015, Fuchs *et al*., 2013, Powell *et al*., 2012 and Powell *et al*., 2015). Intriguingly, it was demonstrated that T-bet^-^ IL-17A^+^ ILC3 are likely to be the drivers of inflammation in this model. In contrast, Rag-sufficient mice with a germline depletion of *Tbx21* have a greater cellularity of intestinal NKp46-negative ILC3 while demonstrating no sign of spontaneous colitis (Schroeder *et al*., 2021). Furthermore, these mice have a greater abundance of ILC2 in colonic lamina propria which may function to limit colitis pathology (Garrido Mesa *et al*., 2019). Hence, we hypothesised that targeted T-bet depletion in ILC has the potential to be a safe approach to treat patients with IBD.

In the current study, we report that in an approach to test the feasibility of targeting T-bet-expressing ILC, temporally induced depletion of T-bet using a Cre-Ert2 model resulted in complete loss of intestinal ILC1 in the lamina propria. In contrast to the loss of these cells, cellularity of ILC2 and NKp46^-^ ILC3 was not altered in the colonic lamina propria upon induced depletion of T-bet. Although the ability of mesenteric lymph node (MLN) and splenic T cells to express T-bet was not affected substantially in these mice, we observed milder pathology in DSS-induced colitis models, pointing to a potentially important pathogenic role of ILC1. The complete loss of intestinal ILC1 upon induced depletion of T-bet was partially reversible. This minimal ILC1 recovery in the intestine may be driven by reseeding of the tissue by bone marrow (BM) ILC1 precursors in the context of DSS-induced colitis. To identify upstream targets controlling T-bet expression in ILC1, we examined STAT1 (*Stat1*^*-/-*^) or STAT4 (*Stat4*^*-/-*^) deficient mice. This revealed that STAT1 and STAT4 were functionally redundant for the development and maintenance of intestinal ILC1. However, prior to the loss of intestinal ILC1, these cells lose IL-15R and NKp46 expression upon induced depletion of T-bet *in vivo*, and these surface markers may therefore function as candidates to support ILC1 maintenance. These data highlight the important role of T-bet in ILC during colitis and a proof-of-concept to support therapeutic targeting of these putatively pathogenic cells.

## RESULTS

### Intestinal lamina propria ILC1 require T-bet for homeostatic tissue maintenance

T-bet is a crucial factor for the development of ILC1 and NKp46^+^ ILC3. However, its continued role in the ongoing peripheral maintenance of these cells has not been addressed. To test whether T-bet is also required for the peripheral maintenance of T-bet^+^ ILC, we created a model allowing for the induced depletion of *Tbx21* in adult mice, using the Rosa26-Cre-Ert2 system (Metzger *et al*., 1995, Schwenk *et al*., 1998). In this model, injections of tamoxifen in the peritoneal cavity are employed temporally to promote the translocation of Cre-Ert2 into the nucleus. Throughout this study ILC were defined as live CD45^+^ Lin^-^ CD127^+^ leukocytes (Supplementary Figure 1a). Colonic lamina propria (cLP) ILC1 were present in untreated *Tbx21*^*fl/fl*^ x *Rosa26-Cre-Ert*^*+/-*^ (*Tbx21*^Δ^) and *Tbx21*^*fl/fl*^ x *Rosa26-Cre-Ert2*^*-/-*^ (*Tbx21*^*fl*^) mice (Supplementary Figure 1b). Tamoxifen was injected into *Tbx21*^Δ^ and *Tbx21*^*fl*^ mice each day for 5 consecutive days (Supplementary Figure 1c). Partial depletion of cLP NKp46^+^ NK1.1^+^ ILC occurred in *Tbx21*^Δ^ mice as early as 1 week after the first injection and strongly reduced numbers of cLP and small intestinal lamina propria (SI LP) NKp46^+^ NK1.1^+^ ILC could be detected 3 weeks after the first injection (Fig 1a, b; Supplementary Figure 1d). NKp46^+^ NK1.1^+^ ILC are a heterogenous population of all ILC1 with a smaller subpopulation of NK1.1^+^ ILC3 (Klose *et al*., 2014). Furthermore, in contrast to colonic and SI lamina propria ILC1, colonic and SI intraepithelial lymphocyte (IEL) ILC1 were less affected by induced depletion of T-bet indicating that this transcription factor is not critical for their maintenance (Figure 1c d). Colonic IEL ILC1 had a reduced cellularity and expression of NKp46 and NK1.1 upon induced depletion of T-bet (Figure 1d-f). These observations were not reflected in SI IEL ILC1, as in these cells only NK1.1 expression was partially dependent on T-bet (Figure 1d-f). While ILC1 were depleted in the colonic and SI lamina propria of treated *Tbx21*^Δ^ mice, NK cells defined as CD49b^+^ NK1.1^+^ CD226^+^ CD3^-^ leukocytes were virtually absent at these sites in *Tbx21*^Δ^ and *Tbx21*^*fl*^ mice (Supplementary Figure 2).

**Figure 1.**
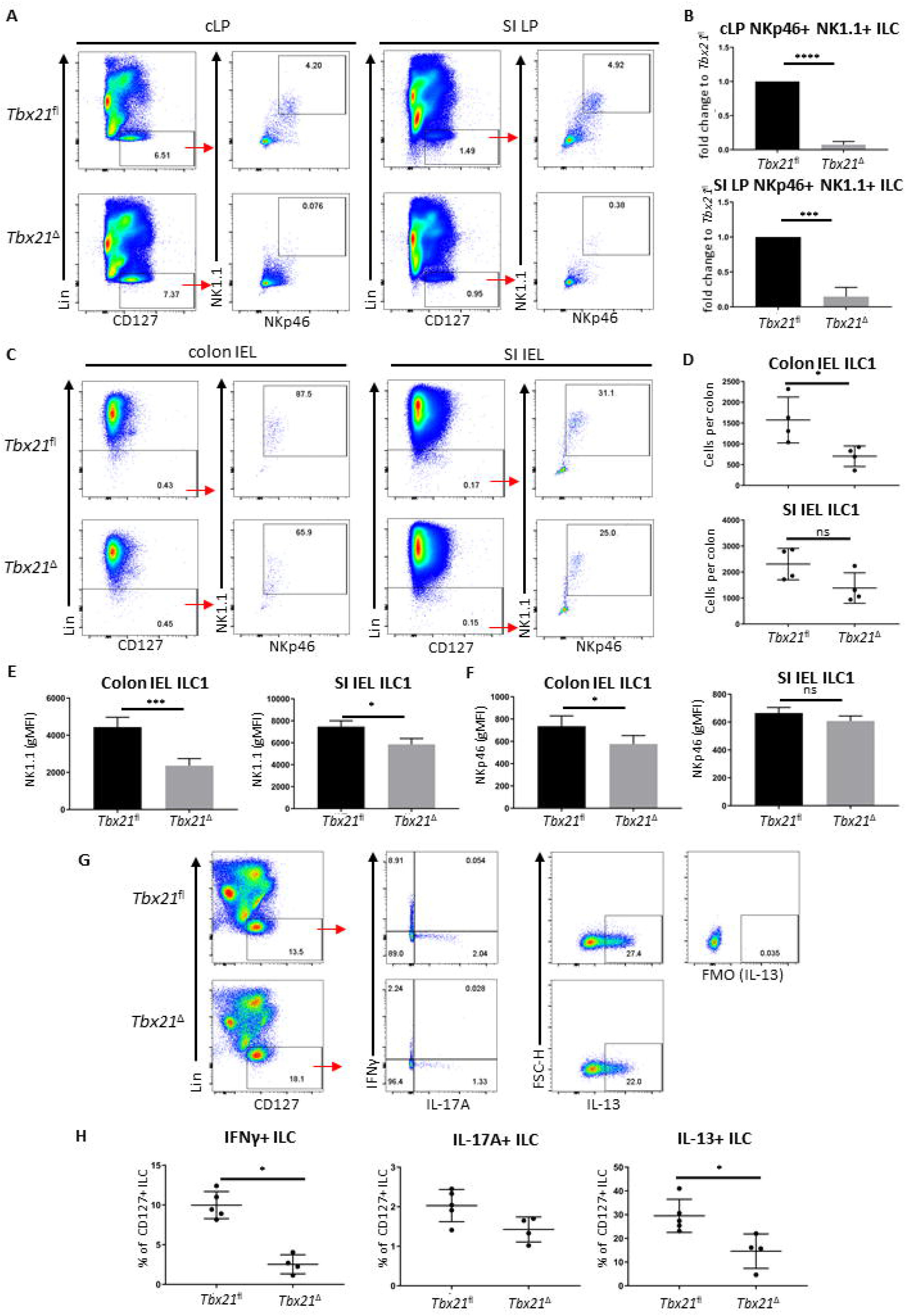
Induced T-bet depletion *in vivo* causes complete loss of intestinal ILC1 ILC were isolated from tamoxifen-treated *Tbx21*^*fl*^ and *Tbx21*^Δ^ mice for flow cytometry analysis 21 days after the first injection of tamoxifen. (A, B) cLP and SI LP ILC1 were analysed as live CD45^+^ Lin^-^ CD127^+^ NKp46^+^ NK1.1^+^ leukocytes and (B) the percentage fold change of NKp46^+^ NK1.1^+^ ILC of total CD127^+^ ILC in the *Tbx21*^Δ^ mice in comparison to the *Tbx21*^*fl*^ controls was determined (n=3). Intestinal IEL ILC were isolated from tamoxifen-treated *Tbx21*^*fl*^ and *Tbx21*^Δ^ mice for flow cytometry analysis. IEL ILC1 were analysed as live CD45^+^ Lin^-^ NKp46^+^ NK1.1^+^ leukocytes. (C, D) Colon and SI IEL ILC1, their cellularity and expression of (E) NK1.1 and (F) NKp46 are shown (n=3). Tamoxifen-pretreated *Tbx21*^*fl*^ and *Tbx21*^Δ^ mice were infected with *C. rodentium*. cLP leukocytes isolated at day 6 post infection were re-stimulated with PMA and ionomycin for 3 hours prior to flow cytometry analysis. (G) Flow cytometry analysis of IFNγ, IL-17A and IL-13 expression by live CD45^+^ Lin^-^ CD127^+^ cLP ILC and (H) statistical analysis of cytokine expression are shown (n=4).

We sought to confirm that the cLP ILC1 depletion observed also resulted in loss of IFNγ-expressing CD127^+^ ILC, which are predominately ILC1. In order to test this, we infected *Tbx21*^Δ^ and *Tbx21*^*fl*^ mice with *Citrobacter rodentium* upon induced deletion of T-bet to establish an inflammatory condition. Induced depletion of T-bet did not cause a profound difference in body weight change over the course of infection (Supplementary Figure 3). However, *Tbx21*^Δ^mice had a substantial loss of IFNγ^+^ ILC supporting the notion that induced depletion of T-bet caused a loss of ILC1. In addition, CD127^+^ ILC did display reduced expression of IL-13, but no altered functionality in terms of IL-17Aproduction upon induced depletion of T-bet (Figure 1e, f).

Further analysis of ILC3 and ILC2 upon induced depletion of T-bet revealed that in contrast to ILC1, the population sizes of these subsets were not significantly altered (Figure 2a-f). For ILC2 analyses, KLRG1 was used as a key marker of intestinal ILC2 in line with recent publications (Flamar *et al*., 2020, Garrido Mesa *et al*., 2019, Schroeder *et al*., 2021). KLRG1 as a marker for intestinal ILC2 has an advantage to GATA3 as intestinal ILC3 have a low expression of GATA3 and the expression of this transcription factor is variable among the ILC2 population (Zhong *et al*., 2016, Zhong *et al*., 2020, Ricardo-Gonzalez *et al*., 2018). KLRG1^hi^ intestinal ILC as gated in this study require GATA3 for post-developmental maintenance supporting the notion these cells are ILC2 (Yagi *et al*., 2014). The expression of RORγt and CD127 in all cLP NKp46-negative ILC3 subsets was unchanged by induced depletion of T-bet, and the expression of RORγt and CD127 were not altered in cLP NKp46^+^ ILC3 (Figure 2g, h). However, cLP NKp46^+^ ILC3 had a lower expression of NKp46, indicating induced depletion of T-bet had some limited effect on these cells (Figure 2i). Overall, there appeared to be no effect of induced depletion of T-bet on cLP NKp46^-^ ILC3 and ILC2 and a limited effect for NKp46^+^ ILC3 in terms of NKp46 expression levels.

**Figure 2.**
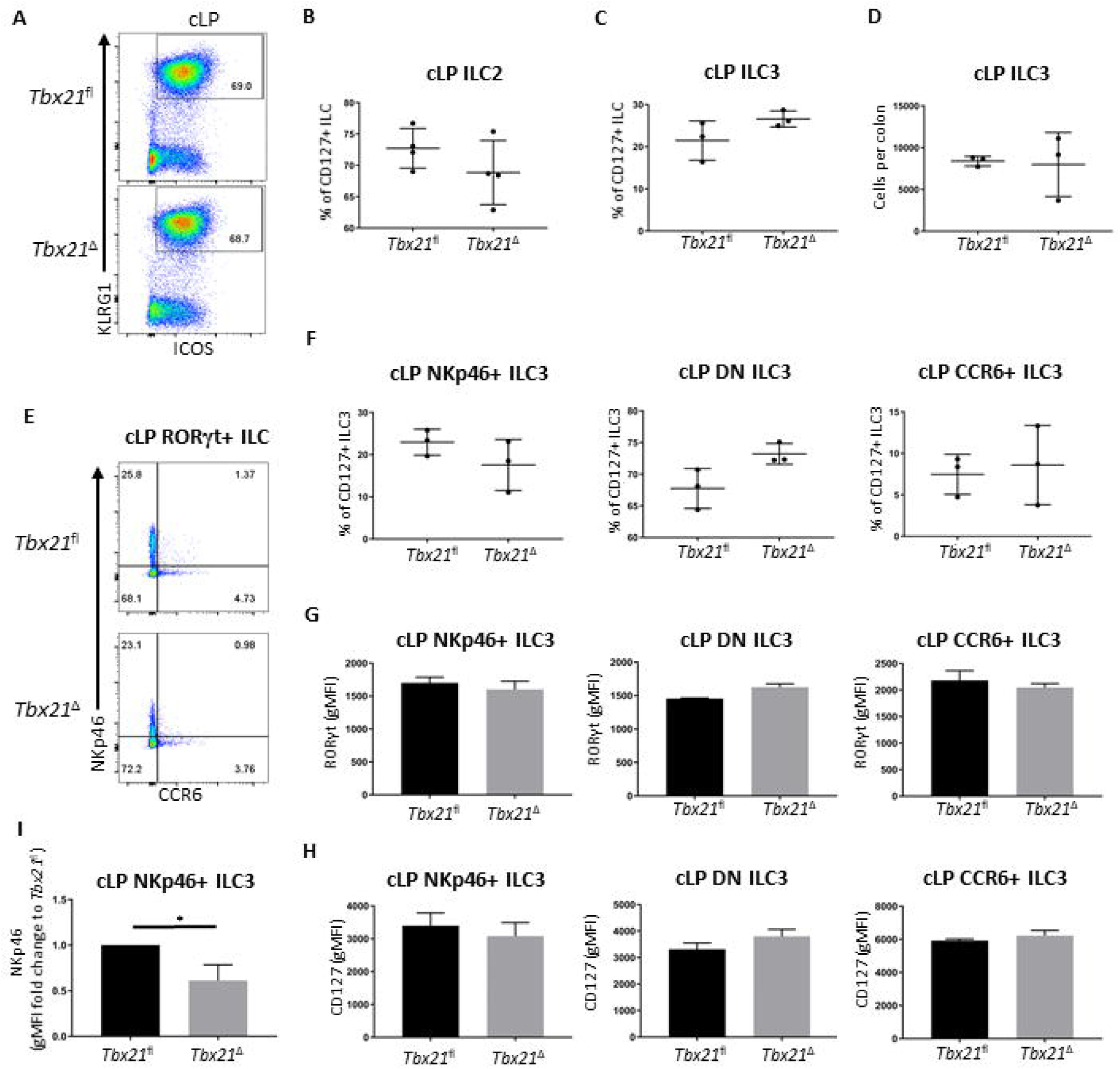
Induced T-bet depletion *in vivo* does not alter cellularity of cLP ILC3 and ILC2 *Tbx21*^*fl*^ and *Tbx21*^Δ^ mice were treated with tamoxifen and flow cytometry analysis on intestinal ILC was performed 21 days after the first injection of tamoxifen. (A) Flow cytometry analyses of cLP ILC2 from *Tbx21*^*fl*^ and *Tbx21*^Δ^ mice and (B) the percentage of cLP ILC2 from total CD127^+^ ILC are shown (n=4). (C) Frequency of RORγt^+^ ILC (ILC3) among CD127^+^ ILC and (D) ILC3 cellularity per colon are demonstrated. (E) cLP NKp46^+^, CCR6^+^ and NKp46^-^ CCR6^-^ double negative (DN) ILC3 were analysed as live CD45^+^ Lin^-^ CD127^+^ RORγt^+^ leukocytes. (F) Percentage of NKp46^+^, DN and CCR6^+^ ILC3 among ILC3 in *Tbx21*^*fl*^ and *Tbx21*^Δ^ mice are outlined (n=3). (G) RORγt and (H) CD127 expression geometric median of fluorescence intensity (gMFI) in NKp46^+^, NKp46^-^ CCR6^-^ DN and CCR6^+^ ILC3 from *Tbx21*^*fl*^ and *Tbx21*^Δ^ mice are illustrated (n=3). (I) gMFI of NKp46 expression fold change expression in in cLP NKp46^+^ ILC3 from *Tbx21*^*fl*^ and *Tbx21*^Δ^ mice are shown (n=3).

### Induced depletion of T-bet results in reduced T-bet expression in ILC1 prior to loss of cell numbers

To further probe a role for T-bet in cLP ILC1 maintenance, these cells were analysed one week prior to the time point at which ILC1 numbers were profoundly reduced in this model (Figure 1a). Two weeks after the first injection of tamoxifen, cLP NKp46^+^ NK1.1^+^ ILC were still detectable in *Tbx21*^Δ^ mice, but their frequency was already significantly reduced (Figure 3a, b). Further analysis revealed that the counts of RORγt^-^ NKp46^+^ cLP ILC1 per colon did not reach a statistically significant loss in comparison to *Tbx21*^*fl*^ cLP ILC1, indicating a gradual decline of ILC1 upon induced depletion of T-bet (Figure 3c, d). Although still present, cLP *Tbx21*^Δ^ ILC1 had a reduced surface expression of CD122, NKp46 and NK1.1 (Figure 3e). Critically, *Tbx21*^Δ^ ILC1 and NKp46^+^ ILC3 showed a significant reduction in T-bet expression, demonstrating T-bet expression was indeed altered at this time point (Figure 3f). cLP NKp46^+^ NK1.1^+^ ILC were present in *Stat1*^*-/-*^ or *Stat4*^*-/-*^ mice, and the percentage of these cells among total CD127^+^ ILC did not alter due to deficiency of either gene (Figure 3g-i). STAT4 deficiency also did not alter T-bet expression in cLP NKp46^+^ NK1.1^+^ ILC (Supplementary Figure 4a, b). The absence of STAT1 or STAT4 did cause significantly altered IFNγ production in CD127^+^ cLP ILC, indicating that STAT1 and STAT4 are critical factors for ILC1 functionality, but redundant for ILC1 maintenance (Figure 3j).

**Figure 3.**
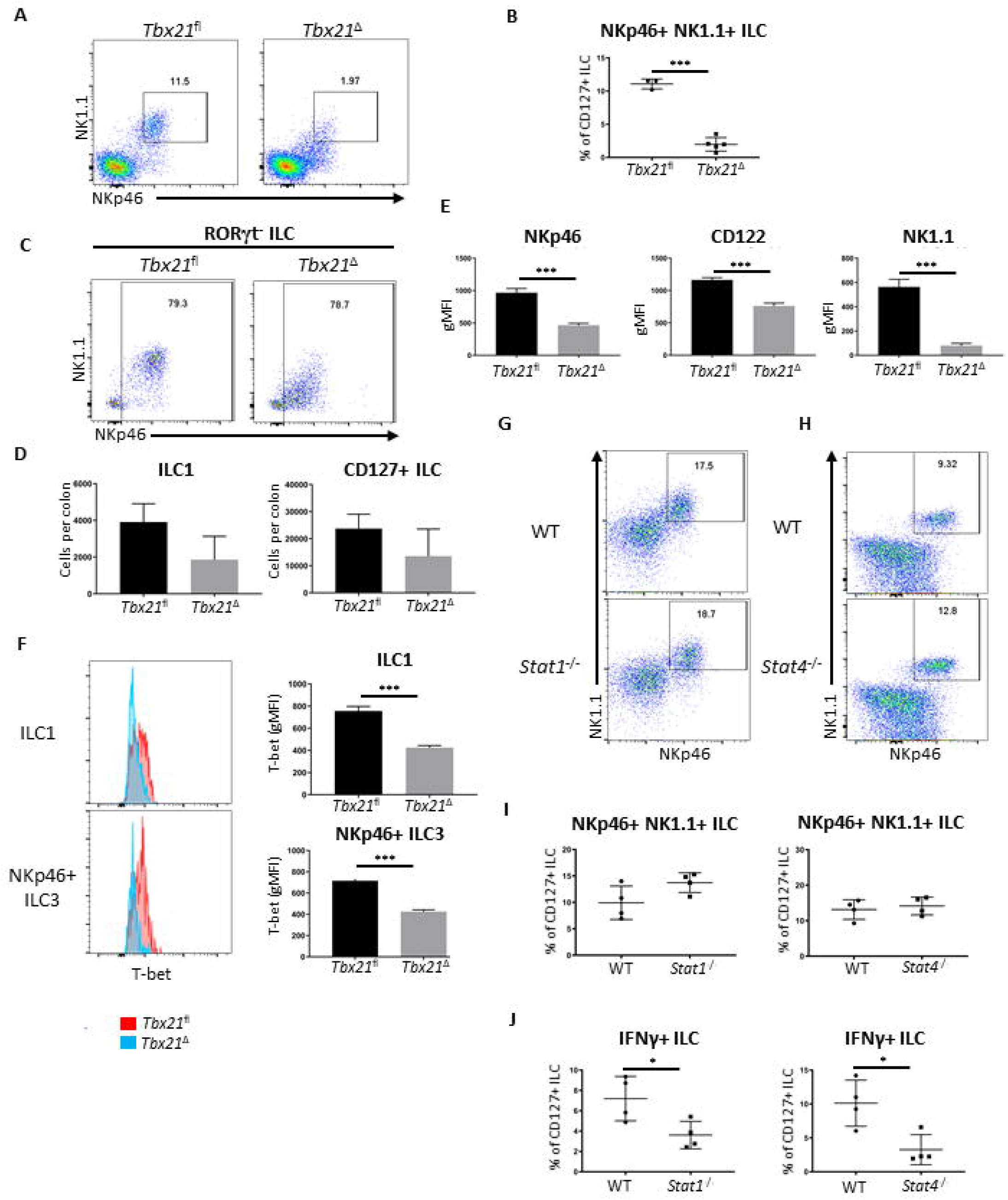
Surface marker expression in ILC1 is altered upon induced deletion of T-bet cLP leukocytes were isolated from tamoxifen-treated *Tbx21*^*fl*^ and *Tbx21*^Δ^ mice for flow cytometry analysis 14 days after the first injection of tamoxifen. (A) Flow cytometry analysis of NKp46^+^ NK1.1^+^ cLP ILC and (B) respective frequency statistical analysis among CD127^+^ ILC are outlined. (C) cLP ILC1 were analysed as CD127^+^ KLRG1^-^ RORγt^-^ NKp46^+^ NK1.1^+/-^ ILC (ILC1) and (D) the cell counts per colon in *Tbx12*^*fl*^ and *Tbx21*^Δ^ mice were determined for ILC1 and CD127^+^ ILC (n=3). (E) gMFI of NKp46, CD122 and NK1.1 surface expression in *Tbx21*^*fl*^ and *Tbx21*^Δ^ ILC1 defined as CD127^+^ KLRG1^-^ RORγt^-^ NKp46^+^ NK1.1^+^ ILC was evaluated statistically (n=3). (F) Flow cytometry and statistical analyses of T-bet protein expression in *Tbx21*^*fl*^ and *Tbx21*^Δ^ cLP ILC1 and NKp46^+^ ILC3 defined as CD127^+^ RORγt^+^ NKp46^+^ ILC are presented (n=3). cLP NKp46^+^ NK1.1^+^ CD127^+^ ILC were analysed in WT, (G) *Stat1*^*-/-*^ or (H) *Stat4*^*-/-*^ mice and (I) the percentage of these cells within the CD127^+^ ILC population was analysed (n=4). (J) IFNγ percentual expression in cLP CD127^+^ ILC from *Stat1*^*-/-*^ and *Stat4*^*-/-*^ mice upon a 4 hour stimulation with PMA and ionomycin is shown (n=4).

### ILC1 have a pathogenic role in DSS-induced colitis

Many immune cells including T cells, ILC, NK cells and myeloid cells can express IFNγ (Castro *et al*., 2018). Hence, we sought to establish an overview of IFNγ-producing cLP cells in naïve mice and mice with DSS-elicited colitis (Eichele *et al*., 2017). *Rorc*^GFP^ mice experienced significant body weight loss upon exposure to 3% DSS in the drinking water compared to controls receiving fresh water (FW) only (Supplementary Figure 5a). IFNγ-producing cLP CD45^+^ leukocytes and ILC1 were detected upon DSS treatment (day 8 of our protocol), but also in naïve mice (Supplementary Figure 5b, c). We next analysed known IFNγ-producing cLP cell types, including CD103^-^ T cells and CD103^+^ resident memory T cells, ILC1, NKp46^+^ ILC3 and NK cells and found populations of all of these cell types to express IFNγ in both FW and DSS treated mice (Supplementary Figure 5d-f). Additional IFNγ producing cells not further characterised were also observed in both treatment groups (Supplementary Figure 5d-f). The frequency of IFNγ-producing ILC1 and NKp46^+^ ILC3 among CD45^+^ leukocytes was greater in control mice in comparison to DSS treated mice (Supplementary Figure 5e, f). In contrast, the frequency of resident memory IFNγ^+^ CD103^+^ T cells increased upon DSS treatment (Supplementary Figure 5e, f). While ILC1 and ILC3 were present in FW and DSS-treated mice, we only detected a substantial abundance of CD11b^+^ Eomes^+^ NK cells in DSS-treated mice, but not in naïve mice (Supplementary Figure 5g-i). The overall frequency of CD103^-^ and resident memory CD103^+^ T cells was similar between naïve and DSS-treated mice (Supplementary Figure 5h, i). These data indicate that ILC1 and NKp46^+^ ILC3 are relevant sources of IFNγ in mice at homeostatic baseline and may therefore become important contributors to the onset of colitis.

To assess the functional effect of T-bet depletion and consequent ILC1 loss, we again utilized the DSS model of colitis. Exposure to 3% DSS in the drinking water caused less severe weight loss in T-bet-deficient (*Tbx21*^*-/-*^) mice in comparison to matched wild type C57BL/6 mice (Figure 4a). BALB/c and BALB/c-background *Tbx21*^*-/-*^ mice were more resistant to DSS-induced colitis as these mice did not experience weight loss in this model (Supplementary Figure 6a). To demonstrate a role of T-bet^+^ innate leukocytes in DSS-induced colitis, BALB/c-background *Rag2*^-/-^ and non-colitic *Rag2*^-/-^x*Tbx21*^-/-^ (TRnUC) mice described previously (Powell *et al*., 2012) were exposed to 3 % or 5 % DSS in the drinking water for 5 days (Supplementary Figure 6b, Figure 4b). While 3% DSS was not sufficient to trigger weight loss independent of the T-bet genotype, 5% DSS induced colitis in those mice, however the phenotype was less severe in T-bet deficient mice (Figure 4b) In contrast, BALB/c-background *Rag2*^-/-^x*γc*^-/-^ and *Rag2*^-/-^x*γc*^-/-^x*Tbx21*^-/-^ exposed to 5% DSS were resistant to colitis and only experienced weight loss after 7-8 days, which was independent of T-bet expression (Figure 4c). *Rag2*^-/-^x*γc*^-/-^ mice lack mature ILC (Roberts *et al*., 2021), thus indicating that these cells drive colitis in the absence of an adaptive immune response.

**Figure 4.**
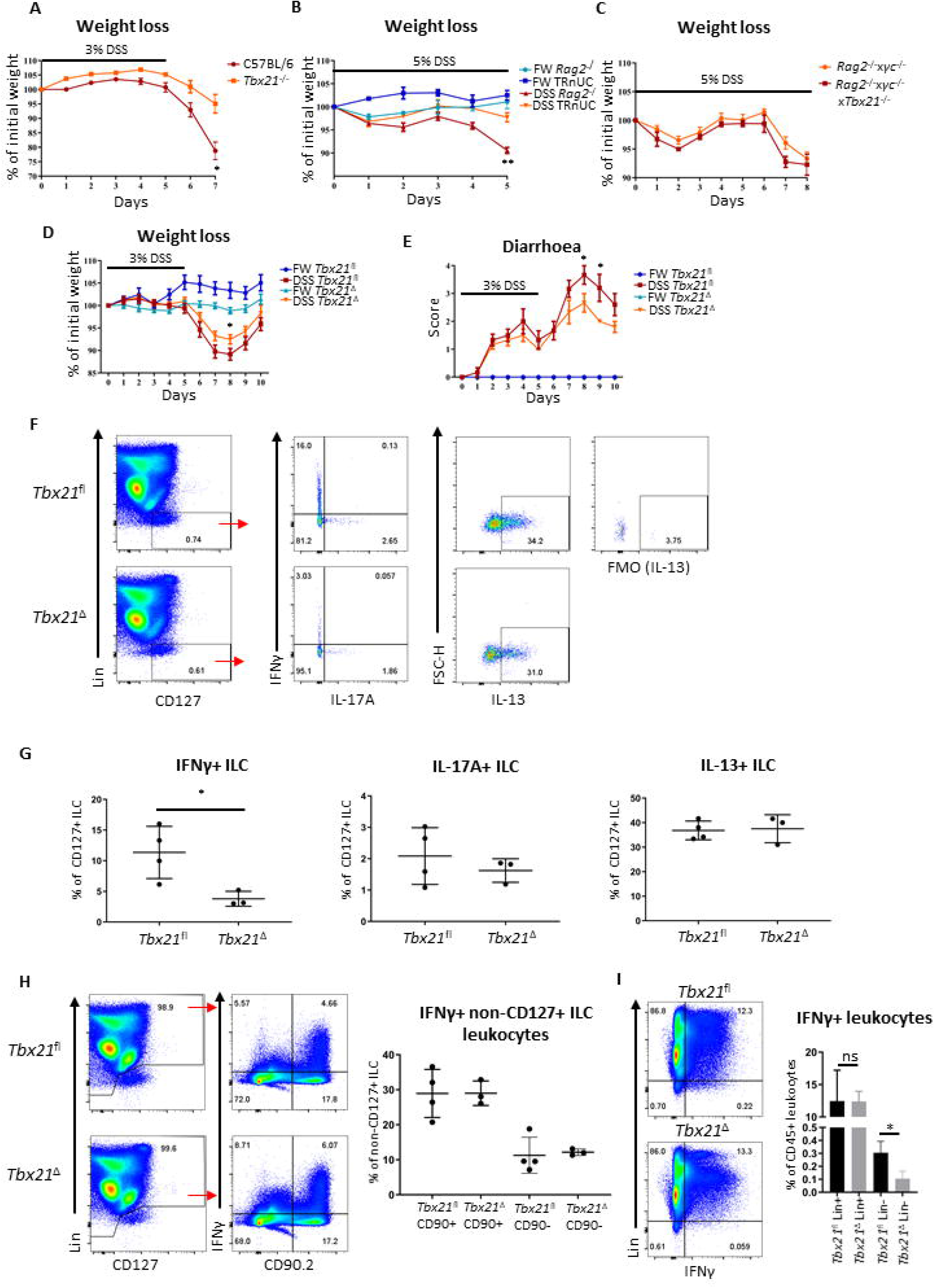
Induced T-bet depletion causes milder DSS-elicited colitis Mice received DSS in the drinking water for weight loss analysis. (A) C57BL/6 and C57BL/6 *Tbx21*^*-/-*^ mice received 3% DSS for 5 days followed by a period of rest (n=4). (B) *Rag2*^-/-^ and TRnUC (n=3) and (C) *Rag*^-/-^x*γc*^-/-^ and *Rag*^-/-^x*γc*^-/-^x*Tbx21*^-/-^ (n=4) received (B) 5% DSS and fresh water (FW) for 5 days followed by a period of rest and (C) 5% DSS for 8 days, respectively. The change in weight in comparison to the starting point are illustrated. (D-E) *Tbx21*^*fl*^ and *Tbx21*^Δ^ mice pre-treated with tamoxifen were exposed to 3% DSS in the drinking water for 5 days followed by a period of rest (n=8-14). Control mice receiving FW without DSS were included in the study. (D) Daily percentual weight change to body weight at the start and (E) daily diarrhoea scoring are illustrated. (F-I) cLP leukocytes were isolated 2 days after DSS withdrawal and re-stimulated with PMA and ionomycin for 3 hours prior to analysis. (F) Flow cytometry analysis of IFNγ, IL-17A and IL-13 expression in live CD45^+^ Lin^-^ CD127^+^ ILC and (G) statistical analysis of cytokine expression are shown (n=3-4). IFNγ expression in (H) CD90.2^+^ and CD90.2^-^ non-ILC leukocytes and (I) live CD45^+^ Lin^-^ and Lin^+^ leukocytes in addition to respective statistical analyses are shown (n=3-4).

To further investigate whether the induced depletion of T-bet also affected the ability of T cells to drive a type 1 immune response, leukocytes from MLN and spleen were isolated from *Tbx21*^*fl*^ and *Tbx21*^Δ^ mice upon induced depletion of T-bet *in vivo* (Supplementary Figure 7a-c). Strikingly, the expression of T-bet in either bulk, naïve or effector CD4^+^ and CD8^+^ T cells was unaltered upon induced depletion of T-bet. Exposure to DSS of tamoxifen-pretreated *Tbx21*^*fl*^ and *Tbx21*^Δ^ mice also did not reveal a significant difference of T-bet expression in either CD4^+^ or CD8^+^ splenic T cells (Supplementary Figure 7d, e). Naïve CD4^+^ T cells were isolated from *Tbx21*^*fl*^ and *Tbx21*^Δ^ MLN and spleen 3 weeks after the first injection of tamoxifen in order to perform an *in vitro* polarization assay. The ability of MLN and splenic naïve CD4^+^ T cells to express T-bet and IFNγ was significantly, but modestly reduced in *Tbx21*^Δ^ T cells in comparison to *Tbx21*^*fl*^ T cells (Supplementary Figure 8a-d). This suggested that T-bet depletion in naive CD4^+^ T cells or T cell precursors was less effective in this mouse model, and T cells escaping gene deletion may occupy tissue niches *in vivo*. These data and the lack of alteration in the ILC2 and ILC3 numbers (Figure 2), suggests that T-bet expression is maintained in many cells in *Tbx21*^Δ^ mice upon temporally induced depletion of the gene encoding this transcription factor. In order to test the relevance of this in an established disease model dependent on intact type 2 immunity, we infected *Tbx21*^Δ^ and *Tbx21*^*fl*^ with *Trichinella spiralis* upon induced depletion of T-bet. We have reported previously that mice with a germline depletion of *Tbx21* are more resistant to an infection with this parasite due to a promoted type 2 immune response (Alcaide *et al*., 2007, Garrido Mesa *et al*., 2019). However, in the model of induced depletion of T-bet, *Tbx21*^Δ^ mice were as resistant to the infection as *Tbx21*^*fl*^ mice with non-significant change in worm burden, crypt and villus length, and muscle thickness (Supplementary Figure 8e, f).

ILC1 have been associated with detrimental outcome of colitis in mice and humans (Bernink *et al*., 2013). To analyse the role of ILC1 in colitis in immunocompetent mice more closely, we tested the course of colitis induced by treatment with DSS in *Tbx21*^Δ^ and *Tbx21*^*fl*^ mice upon induced T-bet depletion. Similar to *Tbx21*^-/-^ mice, *Tbx21*^Δ^ mice showed a milder course of colitis with less weight loss and a lower diarrhoea score than *Tbx21*^*fl*^ mice while mice receiving fresh water did not develop disease (Figure 4d, e). Significant change in body weight at day 8 and diarrhoea score at days 8 and 9 correlated to a significant reduction of IFNγ production in CD127^+^ cLP ILC at day 7 (Figure 4f, g). In contrast to IFNγ, IL-17A and IL-13 production was not different between *Tbx21*^*fl*^ and *Tbx21*^Δ^ CD127^+^ cLP ILC (Figure 4f, g). This indicated again that IFNγ^+^ ILC (being predominately ILC1) were depleted in *Tbx21*^Δ^ mice, while the frequency of *Tbx21*^Δ^ IL-17A^+^ and IL-13^+^ ILC was unaltered upon induced depletion of T-bet. IFNγ can be produced by an abundance of other innate and adaptive leukocytes including NK cells, T cells or neutrophils. However, in contrast to ILC, we did not observe an alteration to IFNγ production in non-ILC CD90^+^ or CD90^−^ leukocytes in the same DSS-treated *Tbx21*^Δ^ mice at the same time point (Figure 4h). A direct comparison of IFNγ expression in lineage marker (Lin)-positive and -negative cLP leukocytes in these mice confirmed that its expression was lower in Lin^-^ cells from *Tbx21*^Δ^ mice in comparison to *Tbx21*^*fl*^ mice while IFNγ production was unaltered in Lin^+^ leukocytes at day 7 (Figure 4i). Hence, these correlative data indicate that ILC1 play a detrimental role during DSS-induced colitis of Rag-sufficient mice.

### DSS colitis promotes limited recolonization of T-bet^+^ ILC in the colonic lamina propria

Targeting of ILC1 to cure colitis may be an attractive therapeutic route, but it would be of relevance to establish whether depletion of cLP ILC1 is reversible. We noticed that *Tbx21*^Δ^ mice exposed to DSS for 5 days 21 days after the initial tamoxifen injection still had fewer cLP NKp46^+^ NK1.1^+^ ILC at day 3 after withdrawal of DSS in comparison to *Tbx21*^*fl*^ mice (Supplementary Figure 9a). However, at this time point, treatment with DSS appeared to have triggered this cell population to re-emerge modestly, as the cellularity was significantly enhanced in comparison to *Tbx21*^Δ^ mice receiving pure fresh water. Exposure to DSS also enhanced the frequency of *Tbx21*^*fl*^ cLP NKp46^+^ NK1.1^+^ ILC among CD127^+^ ILC. Interestingly, cLP NKp46^+^ NK1.1^+^ ILC in *Tbx21*^Δ^ mice were proliferating (or had proliferated recently) at a similar rate to *Tbx21*^*fl*^ cLP NKp46^+^ NK1.1^+^ ILC at day 3 of recovery from DSS- and in FW-treated control mice (Supplementary Figure 9b). Hence, it appeared that upon DSS exposure, the ILC1 population may be re-populated from BM-derived cells rather than as a result of enhanced proliferation of the small number of remaining tissue resident ILC1. This view was further corroborated when we observed that NKp46^+^ NK1.1^+^ ILC were still detectable in the bone marrow upon induced depletion of T-bet (Supplementary Figure 9c). However, the recovered ILC1 cellularity in DSS-treated mice was very minimal and did not reverse their prior loss to a great extent.

## DISCUSSION

There is currently no effective drug to cure IBD, and the need to identify novel strategies of treatment is urgent. In this study we tested T-bet as a treatment target in murine DSS-induced colitis model and report that T-bet and ILC1 depletion in the germline or upon induced depletion in adult mice correlated with less severe DSS-induced colitis.

Induced depletion of T-bet in adult mice resulted in the entire loss of lamina propria ILC1 indicating a critical role for T-bet in post-developmental ILC1 maintenance in the intestine. These data are corroborated by previous reports demonstrating that SI LP ILC1 are virtually absent in *Tbx21*^-/-^ and *NCR1*^iCre^ x *Tbx21*^fl/fl^ mice (Klose *et al*., 2013, Cuff *et al*., 2017). Induced depletion of ILC1 occurred gradually after the onset of the tamoxifen administration regime to deplete T-bet, but once ILC1 were lost, the effect was maintained for several days. A potential and dominant cytotoxic side effect induced by tamoxifen can be excluded as the driver of the depletion as ILC1 in mice lacking the *Cre-Ert2* gene were not affected. cLP NKp46^+^ ILC3 had a significant reduction in NKp46 and loss of NK1.1 surface expression, but were not depleted in mice expressing the *Cre-Ert2* locus. Induced depletion of T-bet did not alter cellularity and functionality of all other ILC3 sub-populations and ILC2 in the colonic lamina propria and overall IL-17A and IL-13 expression from ILC was not affected. This is of critical importance as a pathogenic role of ILC3 has been shown in colitis mouse models with *H. hepaticus* and anti-CD40 antibody driven colitis and even IBD patients (Buonocore *et al*., 2010, Geremia *et al*., 2011, Brasseit *et al*., 2018) and CCR6^+^ ILC3 have been linked to enhanced airway hyperreactivity in an obesity model (Kim *et al*., 2014). However, several studies also highlight the protective functionality of ILC3 in the intestine in DSS colitis and *C. rodentium* infection models (reviewed by Zhou *et al*., 2020).

ILC1 have previously been suggested to play a detrimental role in IBD patients (Bernink *et al*., 2013). ILC1 are also known to mount an early immune response to an infection with MCMV, while the NK and T cell-mediated contribution to host protection is effective later during the course of infection (Weizman *et al*., 2017). We identified innate lymphoid cells as a substantial source of IFNγ at steady state, while other cells like T cells and NK cells dominate more once severe DSS-induced colitis is established. Hence, IFNγ expressing ILC may play a critical role at the onset of colitis. Indeed, when we induced the depletion of T-bet prior to exposure to DSS, there was milder colitis induced by DSS. We did also observe milder DSS-induced colitis in mice with a germline deletion of *Tbx21*, and this was observed in both Rag-sufficient and -deficient mice. In contrast, *Rag2*^-/-^x*γc*^-/-^ mice lacking both mature ILC and adaptive immune response cells were very resistant to developing colitis and the additional germline deletion of T-bet in *Rag2*^-/-^x*γc*^-/-^x*Tbx21*^-/-^ did not change the course of colitis. These results indicated that among innate cells, T-bet^+^ innate lymphocytes play a dominant role in driving colitis. *Tbx21*^-/-^ mice have a non-critical reduction in NK cell development and tissue frequency indicating T-bet is less relevant to maintain the NK cell population in comparison to its essential role for ILC1 (Simonetta *et al*., 2016, Garrido Mesa *et al*., 2019). Considering that the depletion of T-bet also resulted in limited effects in T cells, NKp46^+^ ILC3 and IEL ILC1, we suggest this may indicate that the milder colitis in these models is predominately caused by the lack of lamina propria CD127^+^ ILC1. In fact, splenic and MLN-derived CD4^+^ T cells remained very potent IFNγ producers upon induced depletion of T-bet indicating that the depletion of this transcription factor did not occur in the majority of these cells and overall had no long-term effect in these cell populations. In addition, it is plausible to suggest that Cre-mediated deletion of floxed *Tbx21* requires chromatin remodeling, and the chromatin state of floxed *Tbx21* may control Cre access to the loxP sites. This may be of particular relevance for lymphocyte precursors in the BM and could explain why BM ILC1 and other IFNγ-producing cells like T cells can be found upon induced depletion of T-bet. The difference between ILC1 and Th1 cells for instance could be explained by the greater cell residency of ILC1 while T cells are constitutively generated (Gasteiger *et al*., 2015, O’Sullivan *et al*., 2016, Moro *et al*., 2016, Boulenouar *et al*., 2017). *Tbx21* may be silenced in T cell precursors at the time of tamoxifen exposure leading to a profound T-bet^+^ T cell population despite induced depletion of T-bet. Therefore, it is also important to consider that in this study, all functional experiments were performed no earlier than 17 days after the last injection of tamoxifen. This may provide sufficient time to replace circulating T-bet^+^ T cells that may be depleted in the short-term after induced depletion of T-bet. The chromatin state is a plausible key determinant of the efficiency of *Tbx21* excision. Further corroborating this notion, induced depletion of T-bet had no impact on a subsequent infection with *T. spiralis*, and this stands in contrast to previous work showing that germline depletion of *Tbx21* resulted in a more effective type 2 immune response upon infection with this parasite (Alcaide *et al*., 2007, Garrido Mesa *et al*., 2019). Overall, we postulate that accessibility of the *Tbx21* gene may be the crucial factor driving the depletion success across cell populations.

ILC are known to be predominately resident in mucosal tissues and few of these cells can be found in the periphery. Hence it is no surprise that although entirely depleted initially, an only minor cLP ILC1 population began only to re-emerge in week 5 post the initial injection of tamoxifen. This effect was promoted by the administration of DSS, but appeared not to be the result of a vigorous proliferation of residual cells that may have escaped the depletion. Hence, the colonic lamina propria may be recolonized from ILC precursors from the bone marrow or tissue resident ILC precursors (Björklund *et al*., 2016, Lim *et al*., 2017, Mazzurana *et al*., 2021). Induced deletion of *Tbx21* did not result in the entire depletion of BM NKp46^+^ NK1.1^+^ ILC, and hence these cells may indeed function in the re-population of the colonic lamina propria. Overall, depletion of ILC1 had a beneficial effect in terms of colitis severity, but the caveat of this approach was a limited recovery of ILC1 cellularity, at least acutely.

STAT1 and STAT4 signalling played a significant role in ILC1 functionality, but not maintenance ruling out a non-redundant role of signalling events elicited by IFNγ, IL-12 or IL-27 for ILC1 maintenance, for instance (Dulson *et al*., 2019, Lighvani *et al*., 2001, Krause *et al*., 2006, Hibbert *et al*., 2003, Kamiya *et al*, 2004, Zhu *et al*., 2012). However, induced depletion of T-bet revealed a reduced expression of CD122 (IL-15RB), NKp46 and NK1.1 in cLP ILC1 a week prior to their loss *in vivo*. Previous studies in mice have indicated that NKp46- and IL-15-mediated signalling may play a role for intestinal ILC1 maintenance *in vivo* (Wang *et al*., 2018, Robinette *et al*., 2017, Klose *et al*., 2014). It is therefore plausible to suggest that T-bet-mediated maintenance of ILC1 is at least partially driven by the ability of T-bet to positively regulate the expression of these molecules.

We have demonstrated that *Tbx21*^-/-^ mice on a BALB/c *Rag2*^*-/-*^ background are more susceptible to *H. thyphlonius*-driven colitis in comparison to conventional BALB/c background *Rag2*^*-/-*^ mice pointing to a protective role of T-bet (Garrett *et al*., 2007, Powell *et al*., 2012, Powell *et al*., 2015). We report now that T-bet deficiency is protective from DSS-induced colitis regardless of whether the treated mice have a Rag-sufficient or - deficient background. Hence, T-bet deficiency can cause protection from colitis, but may cause increased pathology upon infection with specific pathogens in the absence of adaptive immune responses, such as regulatory T cells and potentially sIgA, sIgM and sIgG production in the gut (Garrett *et al*., 2007). We suggest that this study provides an important framework to understand more about a potential treatment approach to target intestinal ILC1 based on direct or indirect T-bet-mediated effects. ILC1-derived IFNγ is likely to cause a delay in tissue repair due to vascular barrier disruption and apoptosis of epithelial cells during inflammation (Langer *et al*., 2019, Schuhmann *et al*., 2011, Jowett *et al*., 2021). Despite a strong association of IFNγ with murine models of colitis, the clinical response of fontolizumab, a humanized anti-IFNγ specific antibody, in IBD patients was not detectable although improved inflammatory scores were observed (Ito *et al*., 2006, Langer *et al*., 2019, Brasseit *et al*., 2018, Hommes *et al*., 2006, Reinisch *et al*., 2006, Reinisch *et al*., 2010, Kashani *et al*., 2019). In contrast anti-IL-12p40 antibodies targeting a component shared by IL-12 and IL-23 has been shown to be more effective and safe in IBD clinical trials (Neurath *et al*., 1995, Kashani *et al*., 2019). An approach to inhibit *Tbx21* expression via gene silencing is already in pre-clinical development (Mohamed *et al*., 2016), while alternatively, the feasibility of targeting the transcription factor using a small molecule inhibitor has been demonstrated in principle with the production of such a drug targeting RORγt (Huh *et al*., 2012). A T-bet targeted treatment approach could provide a short time window for adequate IBD treatment such as the correction of the microbiota prior to ILC1 repopulation of the gut (Mishima *et al*., 2020).

## METHODS

### Animals

C57BL/6 WT and BALB/c WT, *Tbx21*^*-/-*^ (C57BL/6 and BALB/c) and BALB/c *Rag2*^-/-^ (all Charles River) mice were sourced commercially. BALB/c *Rag2*^-/-^xγc^-/-^, *Rag2*^-/-^xγc^-/-^ x*Tbx21*^-/-^ and *T-bet*^*fl/fl*^ mice were previously generated by our group. Rosa26-Cre-ERT2 transgenic mice were a kind gift from Dr Thomas Ludwig (Columbia University) and were created using the Cre-ERT2 construct generated by Pierre Chambon (University of Strasbourg). *T-bet*^*fl/fl*^ mice were crossed with Rosa26-Cre-ERT2 mice to generate *Tbx2*^*fl/fl*^ *Rosa26-Cre-Ert2+*^*/-*^ (*Tbx21*^Δ^) and *Tbx2*^*fl/fl*^ *Rosa26-Cre-Ert2*^-/-^ (*Tbx21*^*fl*^) mice. A colony of colitis-free TRnUC mice was generated as described previously (Powell *et al*., 2012). *Stat1*^-/-^ (*B6*.*129P2-Stat1tm1Dlv*; Durbin *et al*., 1996) and *Stat4*^-/-^ (C57BL/6J- *Stat4em3Adiuj/J*, purchased from The Jackson Laboratory) were housed under specific pathogen-free conditions at the Institute of Animal Breeding and Genetics in Vienna according to Federation of European Laboratory Animal Science Associations (FELASA) guidelines. *C57BL/6N* and *C57BL/6J* mice were purchased from Janvier Labs and used as control mice for *Stat1*^-/-^ and *Stat4*^-/-^ mice, respectively. *Rorc*^GFP^ mice were a kind gift of Dr Gérard Eberl (Institut Pasteur, Paris). Age- and sex-matched mice aged 6-12 weeks were used for experiments.

### Tamoxifen induction of *Tbx21* depletion

A tamoxifen stock solution was prepared by dissolving 1g of tamoxifen (free base: MP biomedicals, LLC) in 18 ml 100% ethanol by heating the suspension briefly at 37°C. Prior to injections this solution was diluted at 1:4.56 in sunflower oil and pre-warmed to 37°C. *In vivo* depletion mice was induced by intraperitoneal injections of tamoxifen on five consecutive days with one injection per day (1 mg per injection). Three weeks after the first injection of tamoxifen tissues were isolated or the mice were infected or treated with DSS as outlined below. In separate experiments tissues were harvested one or two weeks after the first injection of tamoxifen.

### Isolation of intestinal leukocytes

cLP and SI LP leukocytes were isolated using a published method (Gronke *et al*., 2017). Briefly, the epithelium of colons or Peyer’s Patch-free SI was removed by incubation in HBSS lacking Mg^2+^ or Ca^2+^ (Invitrogen) supplemented with EDTA and HEPES. The tissue was further digested in HBSS lacking Mg^2+^ or Ca^2+^ supplemented with 2% foetal calf serum (FCS Gold, PAA Laboratories), 0.5 mg/ml collagenase D, 10 μg/ml DNase I and 1.5 mg/ml dispase II (all Roche). The LP lymphocyte-enriched population was harvested from a 40%-80% Percoll (GE Healthcare) gradient interface. For IEL leukocyte isolations, colons and Peyer’s patch-free SI were cut longitudinally, washed and incubated twice in Ca^2+^ Mg^2+^ free HBSS supplemented with 2 mM DTT and 5 mM EDTA for 20 minutes at 37°C and 100 rpm shaking. After each incubation the tissues were vortexed for 10 seconds and the filtered supernatants were subject to Percoll gradient purification prior to further analyses.

### Flow Cytometry

Flow cytometry was performed using a standard protocol. Fc receptor blocking was carried out with anti-CD16/32 specific antibodies. For ILC analyses a lineage cocktail of antibodies specific for CD3, CD45R, CD19, CD11b, TER-119, Gr-1, CD5 and FcεRI was used. For a complete list of the antibodies utilized see Table 1. A FoxP3 staining kit (ebioscience) was used for intracellular staining of transcription factors and cytokines. In case of cytokine analysis, cells were pre-stimulated with 100 ng/ml PMA and 2 µM ionomycin in the presence of 6 µM monensin for 3-4 hours at 37°C 5% CO_2_ prior to flow cytometry analysis. Samples were acquired using an LSRFortessa™ cell analyser (Becton Dickinson, USA) or a Cytoflex LX™ for the data on *Stat*1^-/-^ and *Stat*4^-/-^ mice. All the data were analysed using FlowJo software (Tree Star, USA).

**Table 1 :**
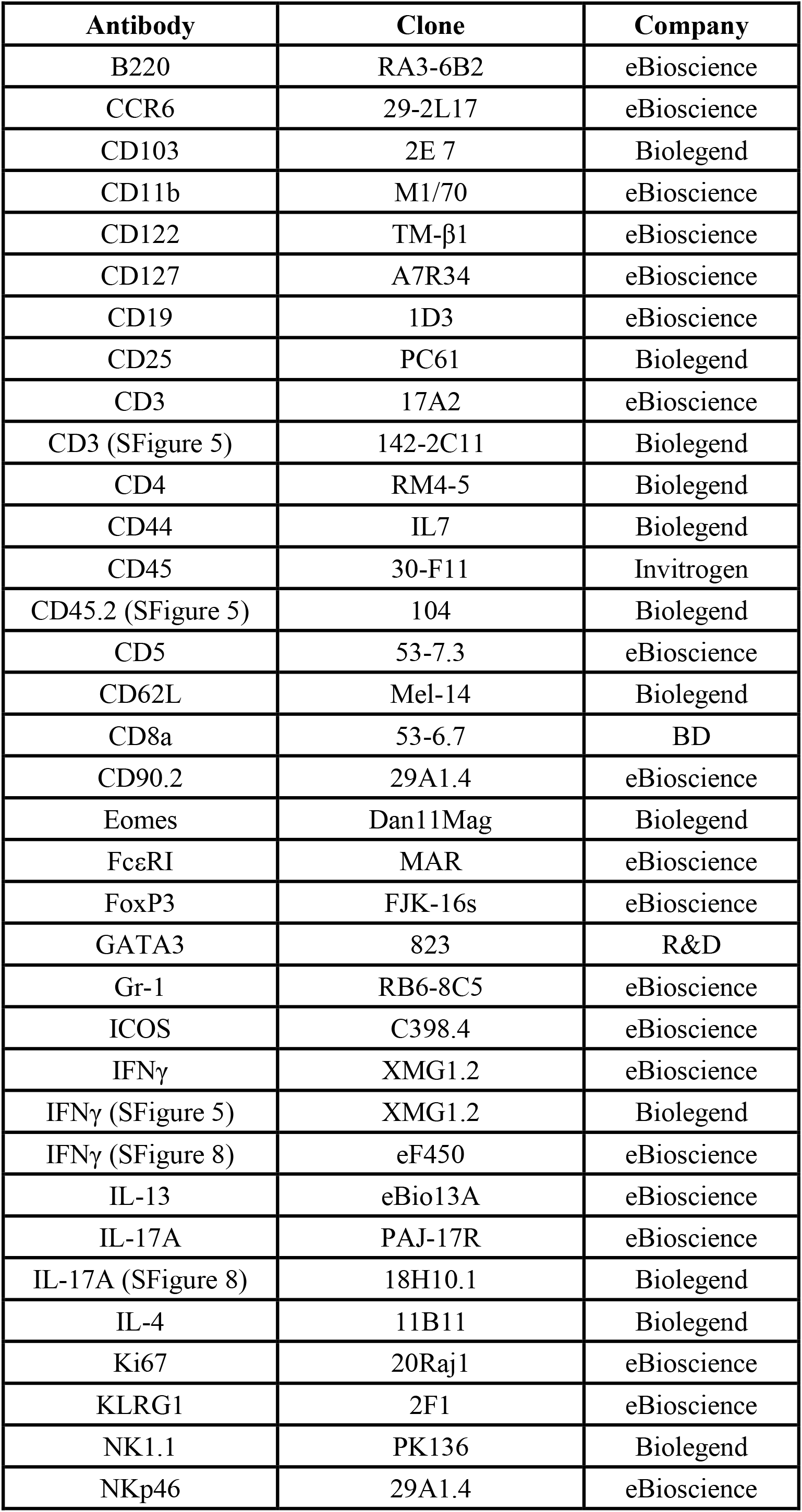
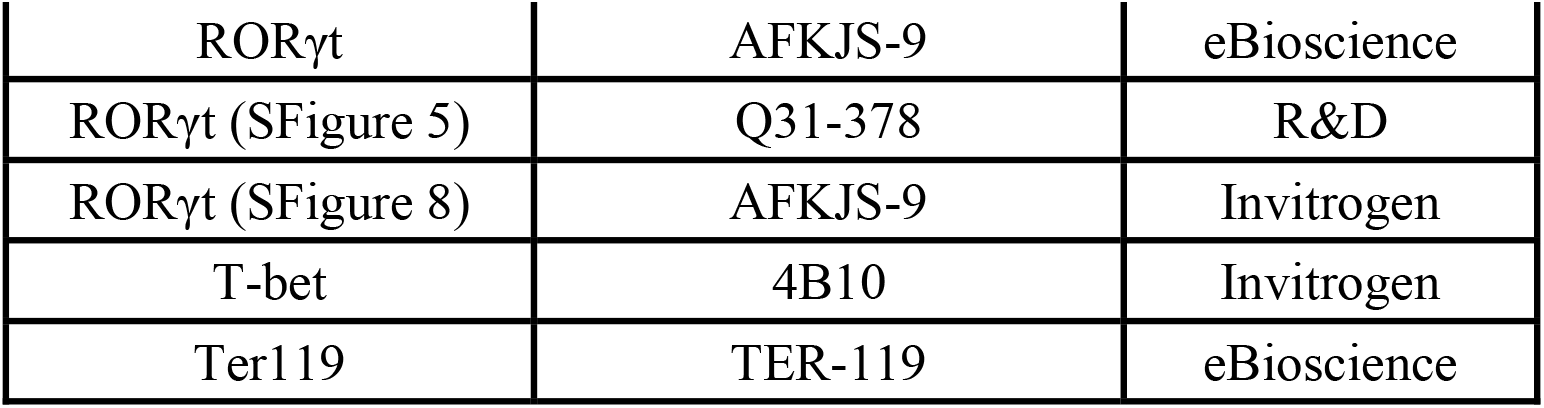
Antibody clones and distributors

### T cell Sorting and polarization assay

Splenocytes and MLN leukocytes were sorted for naïve CD4^+^ cells (live CD4^+^ CD25^-^ CD44^lo^ CD62L^+^) using a BD FACSAria cell sorter (BD Biosciences). 1×10^6^ cells were plated per well of a 48 well plate pre-coated with 2 µg/ml anti-CD3 and anti-CD28 antibodies for 4 hours at 37°C 5% CO_2_. Plates intended to be used for Th17 polarisation were not pre-coated with anti-CD28 antibodies. The sorted cells were cultured in complete RPMI supplemented with either 20 ng/ml rmIL-2 (Th0), 20 ng/ml rhIL-2, 20 ng/ml, rmIL-12 and 5 µg/ml anti-IL4 antibody (Th1), 20 ng/ml IL-2, 20 ng/ml IL-4, 10 µg/ml and anti-IFNγ antibody (Th2) or 1 ng/ml rhTGF-β, 50 ng/ml rmIL-6, 10 ng/ml rmIL-23, 20 µg/ml anti-IFNγ antibody, 10 µg/ml anti-IL-4 antibody and 5 µg/ml anti-CD28 antibody (Th17). After 48 hours in culture, the cells were re-stimulated with the same cytokine and antibody mix. Another 24 hours later the cells were washed and cultured into uncoated 24 well plates in complete RPMI supplemented with the same cytokine and antibody mix. On day 4 of culture, cells were re-stimulated with 100 ng/ml PMA and 2 µM ionomycin in the presence of 6 µM monensin for 4 hours prior to FACS analysis.

### *In vivo* models

Colitis was induced by adding 3% or 5% DSS (36-50 KDa, MP Biomedicals, Ontario, USA) to the drinking water for 5-10 days as indicated. Mice were sacrificed 7-10 days after the beginning of the treatment with DSS. Weight and clinical abnormalities were monitored on a daily basis.

Infection with *C. rodentium* (DSB100) was induced via oral gauvage at a dose of 5×10^8^ CFU per mouse. Prior to the infections, the bacteria were grown in LB Broth to 0.7 OD_600_ and washed into sterile cold PBS.

Mice were infected with *T. spiralis* using a published method (Garrido Mesa *et al*., 2019).

DSS-treated or infected mice were sex- and age-matched and not co-housed with respective control mice.

### Immunohistochemistry

Tissues were fixed in 10% paraformaldehyde and then embedded in paraffin blocks. Microscopically-visible damage in colitis experiments were evaluated by a pathologist observer (KH), who was blinded to the experimental groups.

### Statistics

Results are expressed as mean ± SEM. Data were analysed using Student’s t-test or Mann-Whitney U test, as appropriate, using GraphPad Prism 5.0 (GraphPad Inc., USA). ns: non-significant; *p < 0.05; **p< 0.01; ***p<0.001; ****p<0.0001.

### Study approval

Animal experiments were performed in accredited facilities in accordance with the UK Animals (Scientific Procedures) Act 1986 (Home Office Licence Numbers PPL: 70/6792, 70/8127 and 70/7869). *Stat1*^*-/-*^ and *Stat4*^*-/-*^ mice were bred at the animal facility of the Institute of Animal Breeding and Genetics, University of Veterinary Medicine Vienna according to the guidelines of the Federal Ministry of Science, Research and Economy section 8ff of the Animal Science and Experiments Act, Tierversuchsgesetz [TVG], BMWF-68.205/0068-WF/V/3b/2015.

## ACKNOWLEDGMENTS

We thank the members of the LORD laboratory for valuable discussions and critically commenting on the manuscript. In addition, we thank the BRC FlowCore at King’s College for technical help, Dr Gérard Eberl (Institut Pasteur, Paris) for providing *Rorc*^GFP^ mice and Dr Gitta Stockinger and Dr Andrea Iseppon (Crick Institute, London) for contributing *C. rodentium*.

## AUTHOR CONTRIBUTIONS

Study concept and design (JHS, LBR, JWL, KM, KH, JKH, RKG, JFN, BS, GML), acquisition of data (JHS, LBR, KM, JWL, DH, KH, TZ), data analysis and interpretation (JHS, LBR, JWL, KM, KH, JFN, BS, GML), technical support (ER, CMH, RR), obtained funding (GML, JKH, RKG, BS), drafting of manuscript (JHS), study supervision (GML).

## FIGURE LEGENDS

Supplementary Figure 1. Analysis of cLP ILC1 depletion at days 0 and 7 post onset of induced T-bet depletion

cLP ILC were isolated from untreated or tamoxifen-treated *Tbx21*^*fl*^ and *Tbx21*^Δ^ mice for flow cytometry analysis. (A) ILC were gated as live CD45^+^ Lin^-^ CD127^+^ leukocytes. (B, D) NKp46^+^ NK1.1^+^ ILC (B) from untreated mice (n=5) or (D) 7 days after the first injection of tamoxifen (n=1). (C) Model of induced depletion of T-bet using *Tbx21*^Δ^ and *Tbx21*^*fl*^ control mice.

Supplementary Figure 2. Induced T-bet depletion *in vivo* does not affect intestinal CD49b^+^ NK cells

NK cells were isolated from tamoxifen-treated *Tbx21*^*fl*^ and *Tbx21*^Δ^ mice for flow cytometry analysis 21 days after the first injection of tamoxifen. cLP and SI LP NK cells were analysed as live CD45^+^ CD3^-^ NK1.1^+^ CD226^+^ CD49b^+^ leukocytes (n=1).

Supplementary Figure 3. Induced T-bet depletion has limited effect on colitis caused by *C. rodentium*-infection

Tamoxifen pre-treated *Tbx21*^*fl*^ and *Tbx21*^Δ^ mice were infected with *C. rodentium* 3 weeks after the first injection of tamoxifen for infection analysis. Animal weight difference in relation to day 0 on days post infection are shown. Data shown are representative of 8-9 biological replicates.

Supplementary Figure 4. T-bet-expression in STAT4-deficient ILC

cLP leukocytes were isolated from WT or *Stat4*^*-/-*^ mice. (A) Flow cytometry analysis of T-bet in cLP NKp46^+^ NK1.1^+^ ILC and non-ILC1 cLP ILC and statistical analysis of T-bet expression in cLP NKp46^+^ NK1.1^+^ ILC are illustrated.

Supplementary Figure 5. T-bet-expressing ILC are a relevant source of IFNγ at steady state

RORγt-GFP mice received 3% DSS for 5 days followed by a period of rest. (A) Daily percentual weight change to body weight at the start are illustrated. (B-H) cLP leukocytes were isolated 3 days after DSS withdrawal and re-stimulated with PMA and ionomycin for 3 hours prior to analysis. (B) Flow cytometry analysis of IFNγ expressing live CD45^+^ leukocytes and RORγt^-^ NKp46^+^ NK1.1^+^ ILC1 and (C) respective statistical analyses are shown. Frequency of (D) CD11b^+^ Eomes^+^ NKp46^+^ NK, (E) CD11b^-^ CD103^-^ CD3^+^ T cells, CD11b^-^ CD103^+^ CD3^+^ T cells, RORγt^-^ NKp46^+^ ILC1 and RORγt^+^ NKp46^+^ ILC3 among IFNγ^+^ CD45^+^ leukocytes from DSS-treated and control mice and (F) respective statistical analysis are illustrated. (G) Flow cytometry analysis of RORγt^-^ NKp46^+^ NK1.1^+^ ILC and RORγt^+^ NKp46^+^ ILC3, (H) Eomes^+^ NKp46^+^ NK and CD103^+^ and CD103^-^ T cells and (I) a summary plot of CD45^+^ leukocyte subset frequencies are shown. Data shown are representative of 3 biological replicates.

Supplementary Figure 6. BALB/c *Tbx21*^*-/-*^ mice are resistant to 3% DSS-elicited colitis

Mice received DSS in the drinking water for weight loss analysis. (A) BALB/c and BALB/c *Tbx21*^-/-^ mice and (B) *Rag2*^-/-^ and TRnUC mice received 3% DSS for 5 days followed by a period of rest. Daily recording of weight change is shown. Data shown are representative of 4 biological replicates.

Supplementary Figure 7. Induced T-bet depletion *in vivo* limits, but does not restrict the potency to generate Th1 cells *in vivo*

Leukocytes from MLN and spleen of tamoxifen-treated *Tbx21*^*fl*^ and *Tbx21*^Δ^ mice were isolated for flow cytometry analysis 3 weeks after the first injection. (A) Gating strategy for naïve (CD44^hi^ CD62L^-^) and effector (CD44^low^ CD62L^+^) CD4^+^ and CD8^+^ T cells. (B) Expression of T-bet and (C) statistical expression of its expression in MLN and splenic bulk, naïve and effector CD4^+^ and CD8^+^ T cells are shown (n=3). (D, E) Tamoxifen-pretreated *Tbx21*^*fl*^ and *Tbx21*^Δ^ mice were exposed to 3% DSS for 5 days and splenic leukocytes were isolated another 5 days later for flow cytometry analysis of CD3^+^ T cells. (D) T-bet percentage frequency in CD4^+^ and CD8^+^ T cells and (E) statistical analysis of T-bet expression in these cells are shown (n=4-6).

Supplementary Figure 8. Induced T-bet depletion *in vivo* limits, but does not restrict the potency to generate Th1 cells *in vitro* and does not alter the immune response to *T. spiralis*

T cells were isolated from the spleen of tamoxifen-treated *Tbx21*^*fl*^ and *Tbx21*^Δ^ mice 3 weeks after the first injection for polarization assays and flow cytometry analysis. (A) T-bet, IFNγ, GATA3 and RORγt expression in (A, B) MLN and (C, D) splenic CD4^+^ T cells upon Th0, Th1, Th2 and Th17 polarization is demonstrated. (A, C) Percentage of marker expression and (B, D) T-bet expression are shown (n=4). Tamoxifen pre-treated *Tbx21*^*fl*^ and *Tbx21*^Δ^ mice were infected 3 weeks after the first injection with *T. spiralis* for infection analysis. (E) Histology, (F) worm burden, crypt length, vili length and muscle thickness are shown (n=5).

Supplementary Figure 9. Bone marrow-derived NKp46^+^ ILC can recolonize the intestine upon induced depletion of cLP ILC1

DSS colitis was induced in *Tbx21*^*fl*^ and *Tbx21*^Δ^ mice pre-treated with tamoxifen (i.e. 21 days after the first injection). Mice were exposed to 3% DSS for 5 days and another 3 days later tissues were isolated and CD127^+^ NKp46^+^ NK1.1^+^ ILC were analysed by flow cytometry. In control settings mice received fresh water without DSS. (A) Flow cytometry analysis of cLP NKp46^+^ NK1.1^+^ ILC percentage share of CD127^+^ ILC. (B) Ki67 and T-bet expression in cLP NKp46^+^ NK1.1^+^ ILC and fold change difference of Ki67 expression in NKp46^+^ NK1.1^+^ ILC relating to fresh water-exposed *Tbx21*^*fl*^ mice are illustrated. (C) Cellularity of live CD45^+^ Lin^-^ CD127^+^ NKp46^+^ NK1.1^+^ ILC in the bone marrow at day 3 of DSS withdrawal and the frequency of BM NKp46^+^ NK1.1^+^ ILC within total BM CD127^+^ ILC are demonstrated. Data shown are representative of 3 biological replicates.

## REFERENCES

Alcaide P, Jones TG, Lord GM, Glimcher LH, Hallgren J, Arinobu Y, Akashi K, Paterson AM, Gurish MA, Luscinskas FW. (2007) Dendritic cell expression of the transcription factor T-bet regulates mast cell progenitor homing to mucosal tissue. J Exp Med. 204(2):431–9.

Bal SM, Golebski K, Spits H. (2020) Plasticity of innate lymphoid cell subsets. Nat Rev Immunol. 20(9):552–565.

Bernink JH, Peters CP, Munneke M, te Velde AA, Meijer SL, Weijer K, Hreggvidsdottir HS, Heinsbroek SE, Legrand N, Buskens CJ, Bemelman WA, Mjösberg JM, Spits H. (2013) Human type 1 innate lymphoid cells accumulate in inflamed mucosal tissues. Nat Immunol. 14(3):221–9.

Bernink JH, Krabbendam L, Germar K, de Jong E, Gronke K, Kofoed-Nielsen M, Munneke JM, Hazenberg MD, Villaudy J, Buskens CJ, Bemelman WA, Diefenbach A, Blom B, Spits H. (2015) Interleukin-12 and -23 Control Plasticity of CD127(+) Group 1 and Group 3 Innate Lymphoid Cells in the Intestinal Lamina Propria. Immunity. 43(1):146–60.

Björklund ÅK, Forkel M, Picelli S, Konya V, Theorell J, Friberg D, Sandberg R, Mjösberg J. (2016) The heterogeneity of human CD127(+) innate lymphoid cells revealed by single-cell RNA sequencing. Nat Immunol. 2016 Apr;17(4):451–60.

Boulenouar S, Michelet X, Duquette D, Alvarez D, Hogan AE, Dold C, O’Connor D, Stutte S, Tavakkoli A, Winters D, Exley MA, O’Shea D, Brenner MB, von Andrian U, Lynch L. (2017) Adipose Type One Innate Lymphoid Cells Regulate Macrophage Homeostasis through Targeted Cytotoxicity. Immunity. 46(2):273–286.

Brasseit J, Kwong Chung CKC, Noti M, Zysset D, Hoheisel-Dickgreber N, Genitsch V, Corazza N, Mueller C. (2018) Divergent Roles of Interferon-γ and Innate Lymphoid Cells in Innate and Adaptive Immune Cell-Mediated Intestinal Inflammation. Front Immunol. 9:23.

Buonocore S, Ahern PP, Uhlig HH, Ivanov II, Littman DR, Maloy KJ, Powrie F. (2010) Innate lymphoid cells drive interleukin-23-dependent innate intestinal pathology. Nature. 464(7293):1371–5.

Castro F, Cardoso AP, Gonçalves RM, Serre K, Oliveira MJ. (2018) Interferon-Gamma at the Crossroads of Tumor Immune Surveillance or Evasion. Front Immunol. 9:847

Cuff AO, Male V. (2017) Conventional NK cells and ILC1 are partially ablated in the livers of Ncr1 ^iCre^Tbx21 ^fl/fl^ mice. Version 2. Wellcome Open Res. 2:39.

Dulson SJ, Watkins EE, Crossman DK, Harrington LE. (2019) STAT4 Directs a Protective Innate Lymphoid Cell Response to Gastrointestinal Infection. J Immunol. 203(9):2472–2484.

Durbin JE, Hackenmiller R, Simon MC, Levy DE. (1996) Targeted disruption of the mouse Stat1 gene results in compromised innate immunity to viral disease. Cell. 84(3):443–50.

Eichele DD, Kharbanda KK. (2017) Dextran sodium sulfate colitis murine model: An indispensable tool for advancing our understanding of inflammatory bowel diseases pathogenesis. World J Gastroenterol. 23(33):6016–6029.

Fiancette R, Finlay CM, Willis C, Bevington SL, Soley J, Ng STH, Baker SM, Andrews S, Hepworth MR, Withers DR. (2021) Reciprocal transcription factor networks govern tissue-resident ILC3 subset function and identity. Nat Immunol. In press.

Flamar AL, Klose CSN, Moeller JB, Mahlakõiv T, Bessman NJ, Zhang W, Moriyama S, Stokic-Trtica V, Rankin LC, Putzel GG, Rodewald HR, He Z, Chen L, Lira SA, Karsenty G, Artis D. (2020) Interleukin-33 Induces the Enzyme Tryptophan Hydroxylase 1 to Promote Inflammatory Group 2 Innate Lymphoid Cell-Mediated Immunity. Immunity. 52(4):606-619.e6.

Fuchs A, Vermi W, Lee JS, Lonardi S, Gilfillan S, Newberry RD, Cella M, Colonna M. (2013) Intraepithelial type 1 innate lymphoid cells are a unique subset of IL-12- and IL-15-responsive IFN-γ-producing cells. Immunity. 38(4):769–81.

Garrett WS, Lord GM, Punit S, Lugo-Villarino G, Mazmanian SK, Ito S, Glickman JN, Glimcher LH. (2007) Communicable ulcerative colitis induced by T-bet deficiency in the innate immune system. Cell. 131(1):33–45.

Garrido-Mesa N, Schroeder JH, Stolarczyk E, Gallagher AL, Lo JW, Bailey C, Campbell L, Sexl V, MacDonald TT, Howard JK, Grencis RK, Powell N, Lord GM (2019) T-bet controls intestinal mucosa immune responses via repression of type 2 innate lymphoid cell function. Mucosal Immunol. 12(1):51–63.

Gasteiger G, Fan X, Dikiy S, Lee SY, Rudensky AY. (2015) Tissue residency of innate lymphoid cells in lymphoid and nonlymphoid organs. Science. 350(6263):981–5.

Geremia A, Arancibia-Cárcamo CV, Fleming MP, Rust N, Singh B, Mortensen NJ, Travis SP, Powrie F. (2011) IL-23-responsive innate lymphoid cells are increased in inflammatory bowel disease. J Exp Med. 208(6):1127–33.

Gronke K, Kofoed-Nielsen M, Diefenbach A. (2017) Isolation and Flow Cytometry Analysis of Innate Lymphoid Cells from the Intestinal Lamina Propria. Methods Mol Biol. 1559:255–265.

Hibbert L, Pflanz S, De Waal Malefyt R, Kastelein RA. (2003) IL-27 and IFN-alpha signal via Stat1 and Stat3 and induce T-Bet and IL-12Rbeta2 in naive T cells. J Interferon Cytokine Res. 23(9):513–22.

Hommes DW, Mikhajlova TL, Stoinov S, Stimac D, Vucelic B, Lonovics J, Zákuciová M, D’Haens G, Van Assche G, Ba S, Lee S, Pearce T. (2006) Fontolizumab, a humanised anti-interferon gamma antibody, demonstrates safety and clinical activity in patients with moderate to severe Crohn’s disease. Gut. 55(8):1131–7.

Huh JR, Littman DR. (2012) Small molecule inhibitors of RORγt: targeting Th17 cells and other applications. Eur J Immunol. 42(9):2232–7.

Ishizuka IE, Constantinides MG, Gudjonson H, Bendelac A. (2016) The Innate Lymphoid Cell Precursor. Annu Rev Immunol. 34:299–316.

Ito R, Shin-Ya M, Kishida T, Urano A, Takada R, Sakagami J, Imanishi J, Kita M, Ueda Y, Iwakura Y, Kataoka K, Okanoue T, Mazda O. (2006) Interferon-gamma is causatively involved in experimental inflammatory bowel disease in mice. Clin Exp Immunol. 146(2):330–8.

Jenner RG, Townsend MJ, Jackson I, Sun K, Bouwman RD, Young RA, Glimcher LH, Lord GM. (2009) The transcription factors T-bet and GATA-3 control alternative pathways of T-cell differentiation through a shared set of target genes. Proc Natl Acad Sci U S A. 106(42):17876–81.

Jowett GM, Norman MDA, Yu TTL, Rosell Arévalo P, Hoogland D, Lust ST, Read E, Hamrud E, Walters NJ, Niazi U, Chung MWH, Marciano D, Omer OS, Zabinski T, Danovi D, Lord GM, Hilborn J, Evans ND, Dreiss CA, Bozec L, Oommen OP, Lorenz CD, da Silva RMP, Neves JF, Gentleman E. ILC1 drive intestinal epithelial and matrix remodelling. (2020) Nat Mater. 20(2):250–259.

Kamiya S, Owaki T, Morishima N, Fukai F, Mizuguchi J, Yoshimoto T. (2004) An indispensable role for STAT1 in IL-27-induced T-bet expression but not proliferation of naive CD4+ T cells. J Immunol. 173(6):3871–7.

Karmele EP, Pasricha TS, Ramalingam TR, Thompson RW, Gieseck RL 3rd, Knilans KJ, Hegen M, Farmer M, Jin F, Kleinman A, Hinds DA; 23and Me Research Team, Almeida Pereira T, de Queiroz Prado R, Bing N, Tchistiakova L, Kasaian MT, Wynn TA, Vannella KM. (2019) Anti-IL-13Rα2 therapy promotes recovery in a murine model of inflammatory bowel disease. Mucosal Immunol. 12(5):1174–1186.

Kashani A, Schwartz DA. (2019) The Expanding Role of Anti-IL-12 and/or Anti-IL-23 Antibodies in the Treatment of Inflammatory Bowel Disease. Gastroenterol Hepatol (N Y). 15(5):255–265.

Kim HY, Lee HJ, Chang YJ, Pichavant M, Shore SA, Fitzgerald KA, Iwakura Y, Israel E, Bolger K, Faul J, DeKruyff RH, Umetsu DT. (2014) Interleukin-17-producing innate lymphoid cells and the NLRP3 inflammasome facilitate obesity-associated airway hyperreactivity. Nat Med. 20(1):54–61.

Klose CS, Kiss EA, Schwierzeck V, Ebert K, Hoyler T, d’Hargues Y, Göppert N, Croxford AL, Waisman A, Tanriver Y, Diefenbach A. (2013) A T-bet gradient controls the fate and function of CCR6-RORγt+ innate lymphoid cells. Nature. 494(7436):261–5.

Klose CSN, Flach M, Möhle L, Rogell L, Hoyler T, Ebert K, Fabiunke C, Pfeifer D, Sexl V, Fonseca-Pereira D, Domingues RG, Veiga-Fernandes H, Arnold SJ, Busslinger M, Dunay IR, Tanriver Y, Diefenbach A. (2014) Differentiation of type 1 ILCs from a common progenitor to all helper-like innate lymphoid cell lineages. Cell. 157(2):340–356.

Krause CD, He W, Kotenko S, Pestka S. (2006) Modulation of the activation of Stat1 by the interferon-gamma receptor complex. Cell Res. 16(1):113–23.

Lighvani AA, Frucht DM, Jankovic D, Yamane H, Aliberti J, Hissong BD, Nguyen BV, Gadina M, Sher A, Paul WE, O’Shea JJ. (2001) T-bet is rapidly induced by interferon-gamma in lymphoid and myeloid cells. Proc Natl Acad Sci U S A. 98(26):15137–42.

Langer V, Vivi E, Regensburger D, Winkler TH, Waldner MJ, Rath T, Schmid B, Skottke L, Lee S, Jeon NL, Wohlfahrt T, Kramer V, Tripal P, Schumann M, Kersting S, Handtrack C, Geppert CI, Suchowski K, Adams RH, Becker C, Ramming A, Naschberger E, Britzen-Laurent N, Stürzl M. (2019) IFN-γ drives inflammatory bowel disease pathogenesis through VE-cadherin-directed vascular barrier disruption. J Clin Invest. 129(11):4691–4707.

Lazarevic V, Chen X, Shim JH, Hwang ES, Jang E, Bolm AN, Oukka M, Kuchroo VK, Glimcher LH. (2011) T-bet represses T(H)17 differentiation by preventing Runx1-mediated activation of the gene encoding RORγt. Nat Immunol. 12(1):96–104.

Lim AI, Li Y, Lopez-Lastra S, Stadhouders R, Paul F, Casrouge A, Serafini N, Puel A, Bustamante J, Surace L, Masse-Ranson G, David E, Strick-Marchand H, Le Bourhis L, Cocchi R, Topazio D, Graziano P, Muscarella LA, Rogge L, Norel X, Sallenave JM, Allez M, Graf T, Hendriks RW, Casanova JL, Amit I, Yssel H, Di Santo JP. (2017) Systemic Human ILC Precursors Provide a Substrate for Tissue ILC Differentiation. Cell. 168(6):1086-1100.e10.

Mazzurana L, Czarnewski P, Jonsson V, Wigge L, Ringnér M, Williams TC, Ravindran A, Björklund ÅK, Säfholm J, Nilsson G, Dahlén SE, Orre AC, Al-Ameri M, Höög C, Hedin C, Szczegielniak S, Almer S, Mjösberg J. (2021) Tissue-specific transcriptional imprinting and heterogeneity in human innate lymphoid cells revealed by full-length single-cell RNA-sequencing. Cell Res. 31(5):554–568.

Metzger D, Clifford J, Chiba H, Chambon P. (1995) Conditional site-specific recombination in mammalian cells using a ligand-dependent chimeric Cre recombinase. Proc Natl Acad Sci U S A. 92(15):6991–5.

Mishima Y, Sartor RB. (2020) Manipulating resident microbiota to enhance regulatory immune function to treat inflammatory bowel diseases. J Gastroenterol. 55(1):4–14.

Mohamed R, Lord GM. (2016) T-bet as a key regulator of mucosal immunity. Immunology. 147(4):367–76.

Moro K, Kabata H, Tanabe M, Koga S, Takeno N, Mochizuki M, Fukunaga K, Asano K, Betsuyaku T, Koyasu S. Interferon and IL-27 antagonize the function of group 2 innate lymphoid cells and type 2 innate immune responses. (2016) Nat Immunol. 17(1):76–86.

Neurath MF, Fuss I, Kelsall BL, Stüber E, Strober W.J (1995) Antibodies to interleukin 12 abrogate established experimental colitis in mice. Exp Med. 182(5):1281–90.

O’Sullivan TE, Rapp M, Fan X, Weizman OE, Bhardwaj P, Adams NM, Walzer T, Dannenberg AJ, Sun JC. (2016) Adipose-Resident Group 1 Innate Lymphoid Cells Promote Obesity-Associated Insulin Resistance. Immunity. 45(2):428–41.

Panda SK, Colonna M. (2019) Innate Lymphoid Cells in Mucosal Immunity. Front Immunol. 10:861.

Powell N, Walker AW, Stolarczyk E, Canavan JB, Gökmen MR, Marks E, Jackson I, Hashim A, Curtis MA, Jenner RG, Howard JK, Parkhill J, MacDonald TT, Lord GM. (2012) The transcription factor T-bet regulates intestinal inflammation mediated by interleukin-7 receptor+ innate lymphoid cells. Immunity. 37(4):674–84.

Powell N, Lo JW, Biancheri P, Vossenkämper A, Pantazi E, Walker AW, Stolarczyk E, Ammoscato F, Goldberg R, Scott P, Canavan JB, Perucha E, Garrido-Mesa N, Irving PM, Sanderson JD, Hayee B, Howard JK, Parkhill J, MacDonald TT, Lord GM. (2015) Interleukin 6 Increases Production of Cytokines by Colonic Innate Lymphoid Cells in Mice and Patients With Chronic Intestinal Inflammation. Gastroenterology. 149(2):456-67.e15.

Reinisch W, Hommes DW, Van Assche G, Colombel JF, Gendre JP, Oldenburg B, Teml A, Geboes K, Ding H, Zhang L, Tang M, Cheng M, van Deventer SJ, Rutgeerts P, Pearce T. (2006) A dose escalating, placebo controlled, double blind, single dose and multidose, safety and tolerability study of fontolizumab, a humanised anti-interferon gamma antibody, in patients with moderate to severe Crohn’s disease. Gut. 55(8):1138–44.

Reinisch W, de Villiers W, Bene L, Simon L, Rácz I, Katz S, Altorjay I, Feagan B, Riff D, Bernstein CN, Hommes D, Rutgeerts P, Cortot A, Gaspari M, Cheng M, Pearce T, Sands BE. (2010) Fontolizumab in moderate to severe Crohn’s disease: a phase 2, randomized, double-blind, placebo-controlled, multiple-dose study. Inflamm Bowel Dis. 16(2):233–42.

Ricardo-Gonzalez RR, Van Dyken SJ, Schneider C, Lee J, Nussbaum JC, Liang HE, Vaka D, Eckalbar WL, Molofsky AB, Erle DJ, Locksley RM. (2018) Tissue signals imprint ILC2 identity with anticipatory function. Nat Immunol. 19(10):1093–1099.

Robinette ML, Bando JK, Song W, Ulland TK, Gilfillan S, Colonna M. (2017) IL-15 sustains IL-7R-independent ILC2 and ILC3 development. Nat Commun. 8:14601.

Roberts LB, Jowett GM, Read E, Zabinski T, Berkachy R, Selkirk ME, Jackson I, Niazi U, Anandagoda N, Araki M, Araki K, Kasturiarachchi J, James C, Enver T, Nimmo R, Reis R, Howard JK, Neves JF, Lord GM. (2021) MicroRNA-142 Critically Regulates Group 2 Innate Lymphoid Cell Homeostasis and Function. J Immunol. 206(11):2725–2739.

Schroeder JH, Meissl K, Hromadová D, Lo JW, Neves JF, Howard JK, Helmby H, Powell N, Strobl B, Lord GM. (2021) T-Bet Controls Cellularity of Intestinal Group 3 Innate Lymphoid Cells. Front Immunol. (2021) 11:623324.

Schuhmann D, Godoy P, Weiss C, Gerloff A, Singer MV, Dooley S, Böcker U. (2011) Interfering with interferon-γ signalling in intestinal epithelial cells: selective inhibition of apoptosis-maintained secretion of anti-inflammatory interleukin-18 binding protein. Clin Exp Immunol. 163(1):65–76.

Schwenk F, Kuhn R, Angrand PO, Rajewsky K, Stewart AF. (1998) Temporally and spatially regulated somatic mutagenesis in mice. Nucleic Acids Res. 26(6):1427–32.

Simonetta F, Pradier A, Roosnek E. (2016) T-bet and Eomesodermin in NK Cell Development, Maturation, and Function. Front Immunol. 7:241.

Soderquest K, Hertweck A, Giambartolomei C, Henderson S, Mohamed R, Goldberg R, Perucha E, Franke L, Herrero J, Plagnol V, Jenner RG, Lord GM. (2017) Genetic variants alter T-bet binding and gene expression in mucosal inflammatory disease. PLoS Genet. 13(2): e1006587.

Vonarbourg C, Mortha A, Bui VL, Hernandez PP, Kiss EA, Hoyler T, Flach M, Bengsch B, Thimme R, Hölscher C, Hönig M, Pannicke U, Schwarz K, Ware CF, Finke D, Diefenbach A. (2010) Regulated expression of nuclear receptor RORγt confers distinct functional fates to NK cell receptor-expressing RORγt(+) innate lymphocytes. Immunity. 33(5):736–51.

Wang Y, Dong W, Zhang Y, Caligiuri MA, Yu J. (2018) Dependence of innate lymphoid cell 1 development on NKp46. PLoS Biol. 16(4):e2004867.

Weizman OE, Adams NM, Schuster IS, Krishna C, Pritykin Y, Lau C, Degli-Esposti MA, Leslie CS, Sun JC, O’Sullivan TE. (2017) ILC1 Confer Early Host Protection at Initial Sites of Viral Infection. Cell. 171(4):795-808.e12.

Withers DR, Hepworth MR, Wang X, Mackley EC, Halford EE, Dutton EE, Marriott CL, Brucklacher-Waldert V, Veldhoen M, Kelsen J, Baldassano RN, Sonnenberg GF. (2016) Transient inhibition of ROR-γt therapeutically limits intestinal inflammation by reducing TH17 cells and preserving group 3 innate lymphoid cells. Nat Med. 22(3):319–23.

Yagi R, Zhong C, Northrup DL, Yu F, Bouladoux N, Spencer S, Hu G, Barron L, Sharma S, Nakayama T, Belkaid Y, Zhao K, Zhu J. (2014) The transcription factor GATA3 is critical for the development of all IL-7Rα-expressing innate lymphoid cells. Immunity. 40(3):378–88.

Zhong C, Cui K, Wilhelm C, Hu G, Mao K, Belkaid Y, Zhao K, Zhu J. (2016) Group 3 innate lymphoid cells continuously require the transcription factor GATA-3 after commitment. Nat Immunol. 17(2):169–78.

Zhong C, Zheng M, Cui K, Martins AJ, Hu G, Li D, Tessarollo L, Kozlov S, Keller JR, Tsang JS, Zhao K, Zhu J. (2020) Differential Expression of the Transcription Factor GATA3 Specifies Lineage and Functions of Innate Lymphoid Cells. Immunity. 52(1):83-95.e4.

Zhou W, Sonnenberg GF. (2020) Activation and Suppression of Group 3 Innate Lymphoid Cells in the Gut. Trends Immunol. 41(8):721–733.

Zhu J, Jankovic D, Oler AJ, Wei G, Sharma S, Hu G, Guo L, Yagi R, Yamane H, Punkosdy G, Feigenbaum L, Zhao K, Paul WE. Immunity. (2012) The transcription factor T-bet is induced by multiple pathways and prevents an endogenous Th2 cell program during Th1 cell responses. 37(4):660–73.

