## Supplementary figures and images for "Sustained post-developmental T-bet expression is critical for the maintenance of type one innate lymphoid cells *in vivo*"

### Supplementary Figure 1

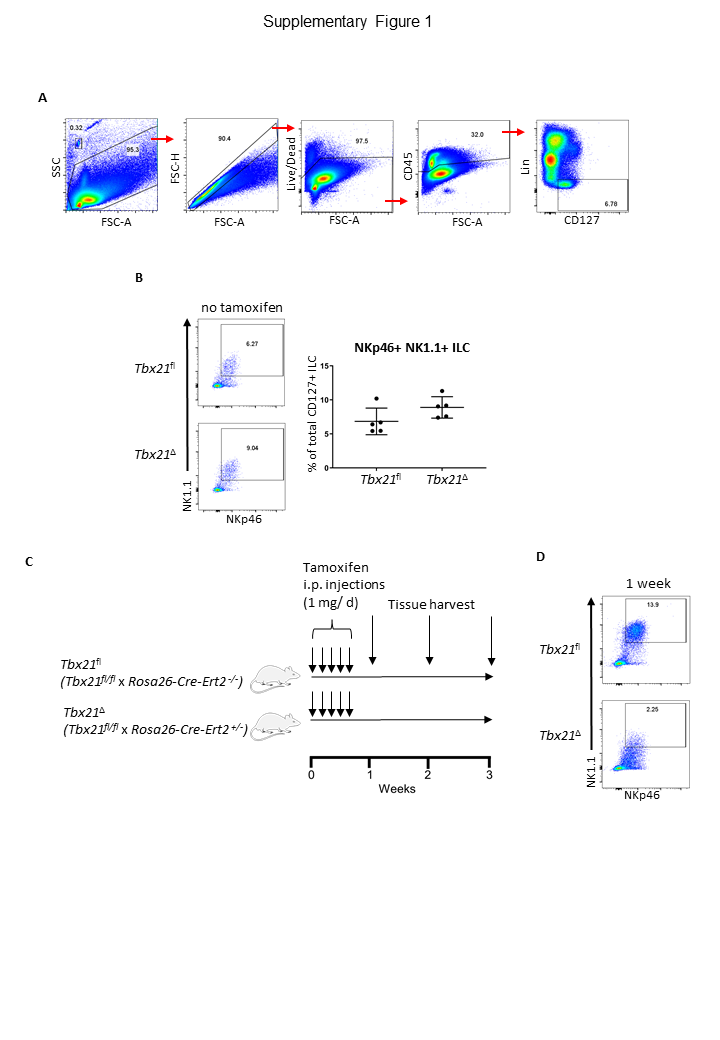

### Supplementary Figure 2

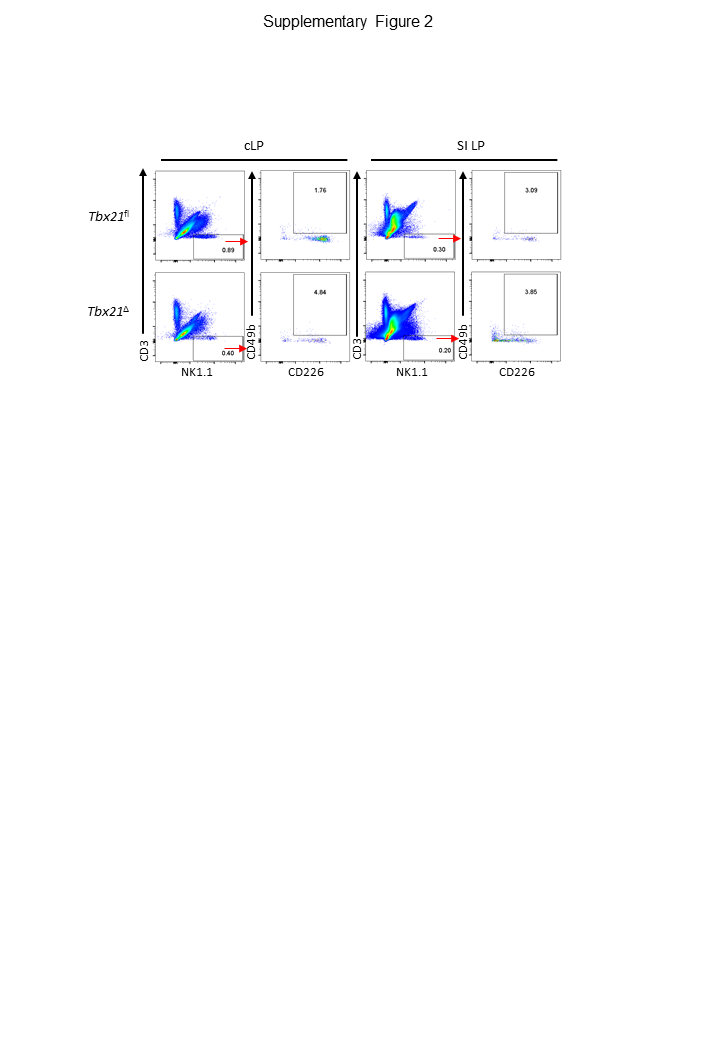

### Supplementary Figure 3

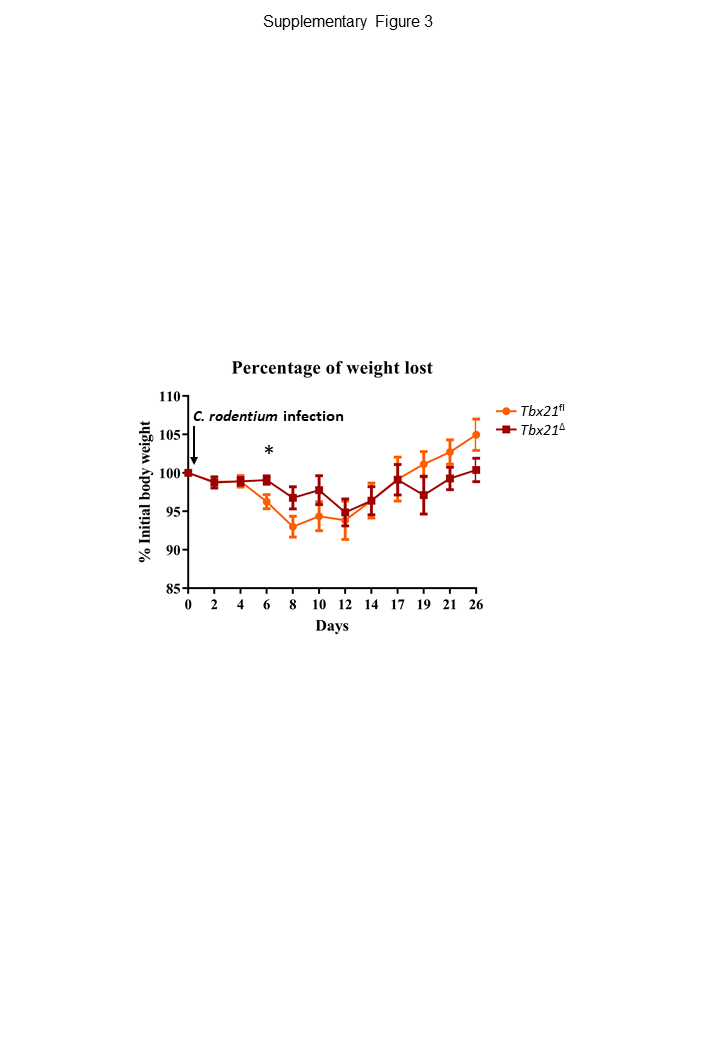

### Supplementary Figure 4

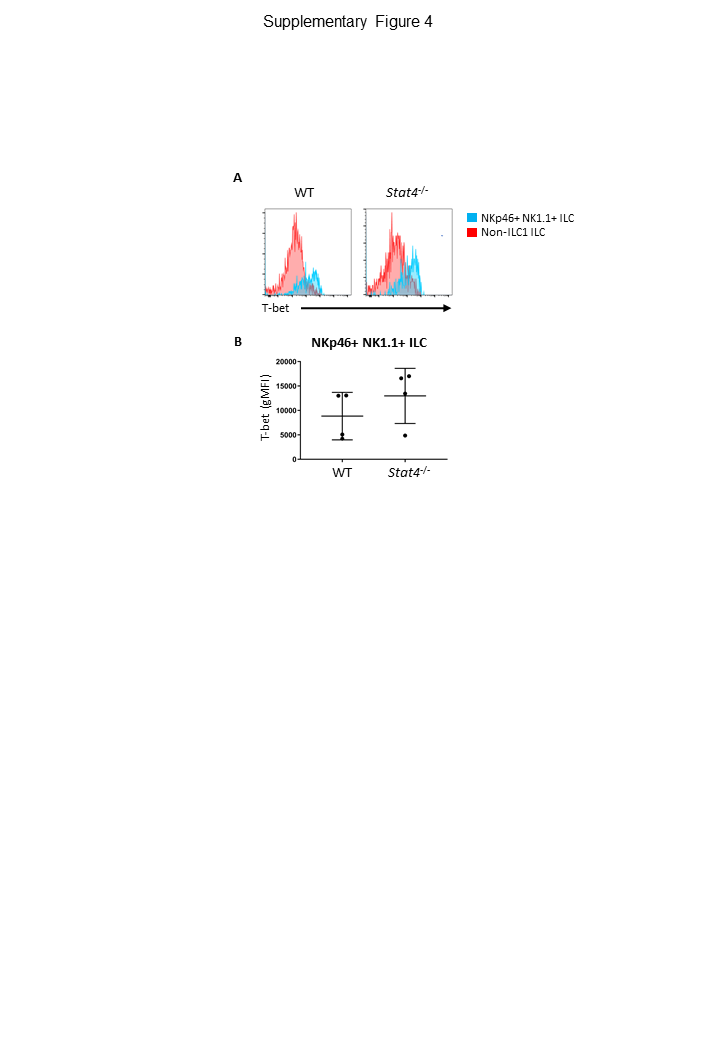

### Supplementary Figure 5

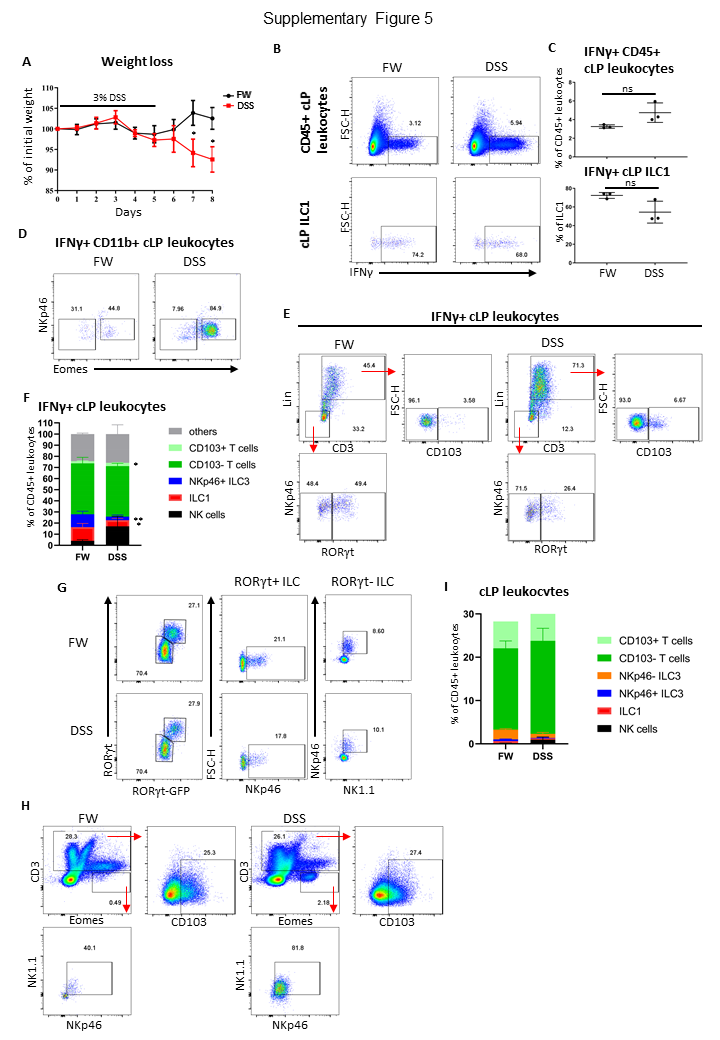

### Supplementary Figure 6

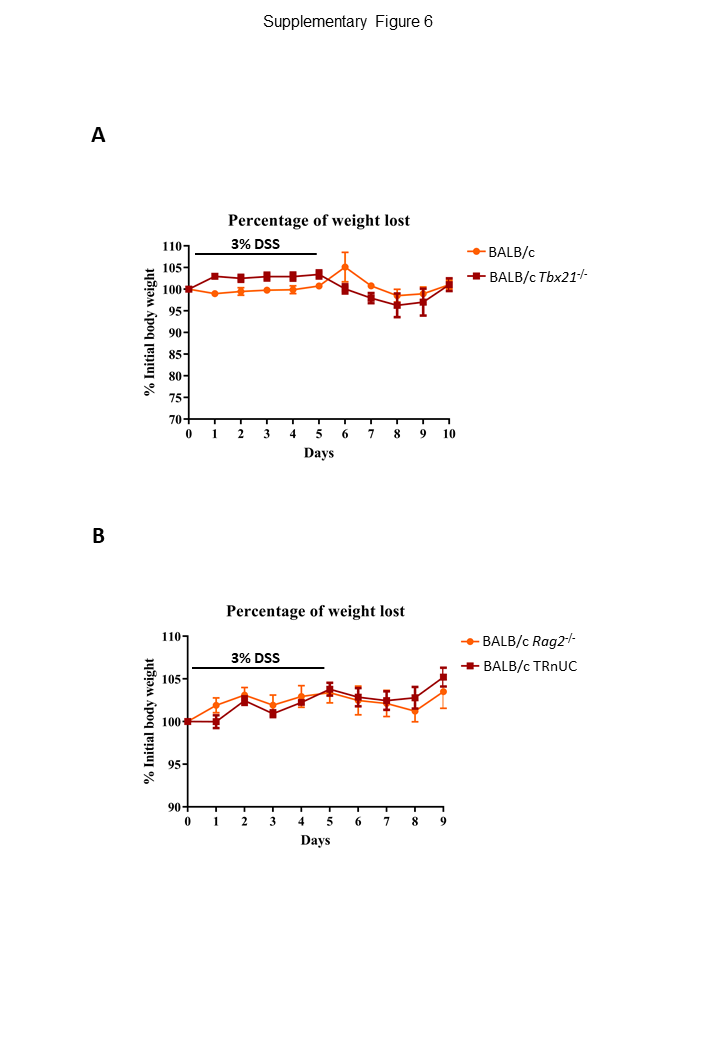

### Supplementary Figure 7

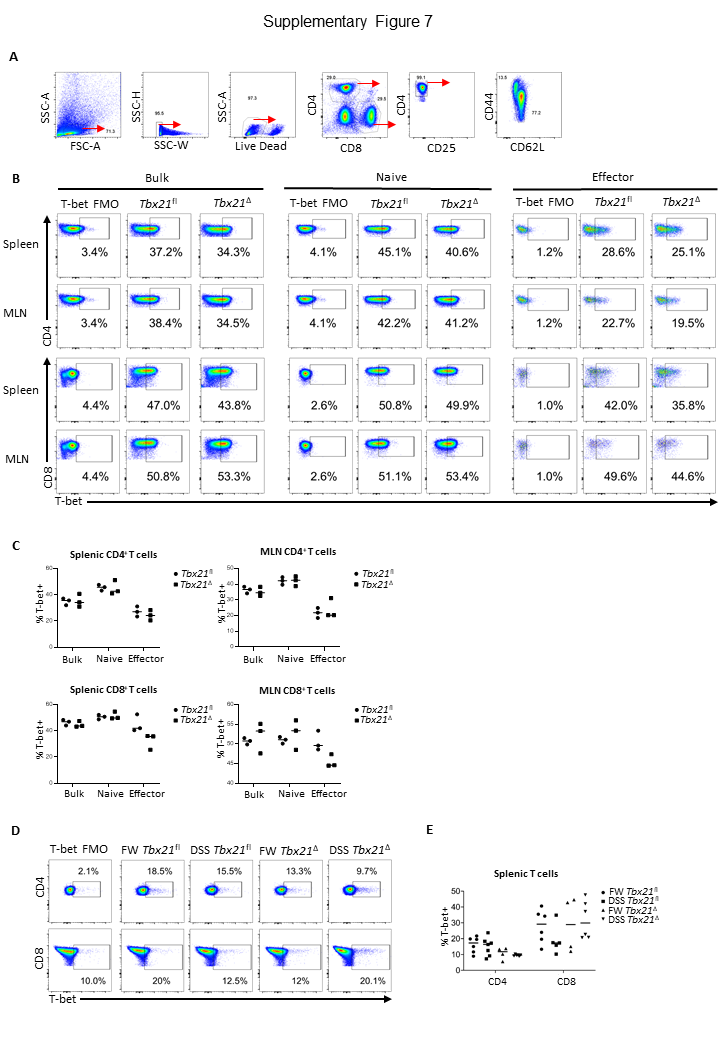

### Supplementary Figure 8

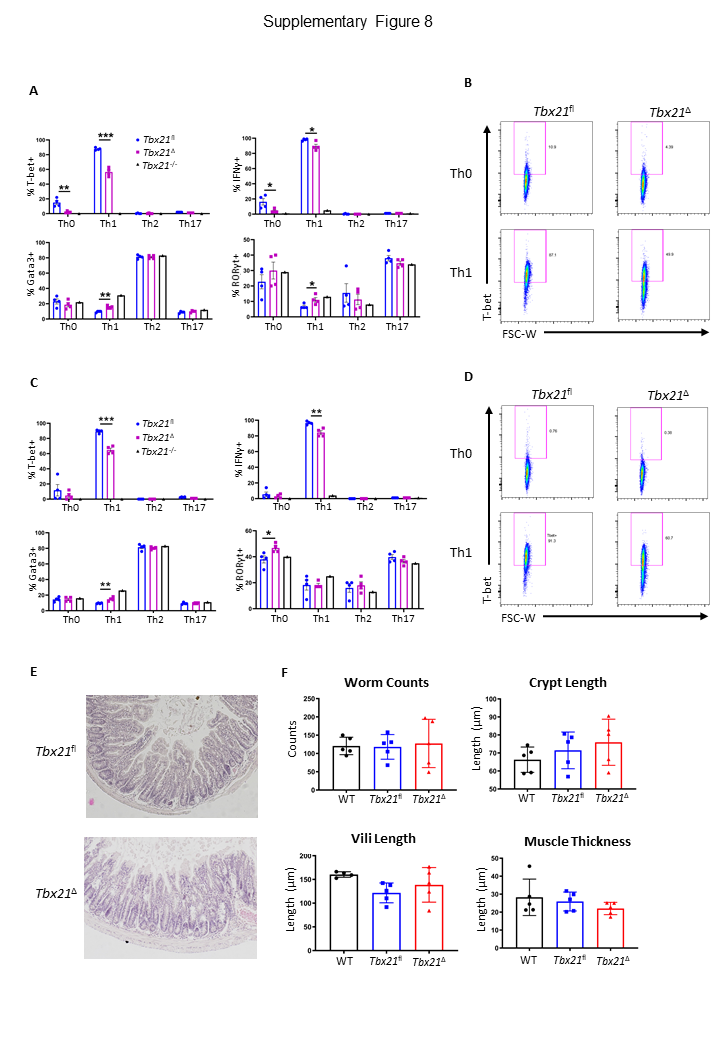

### Supplementary Figure 9

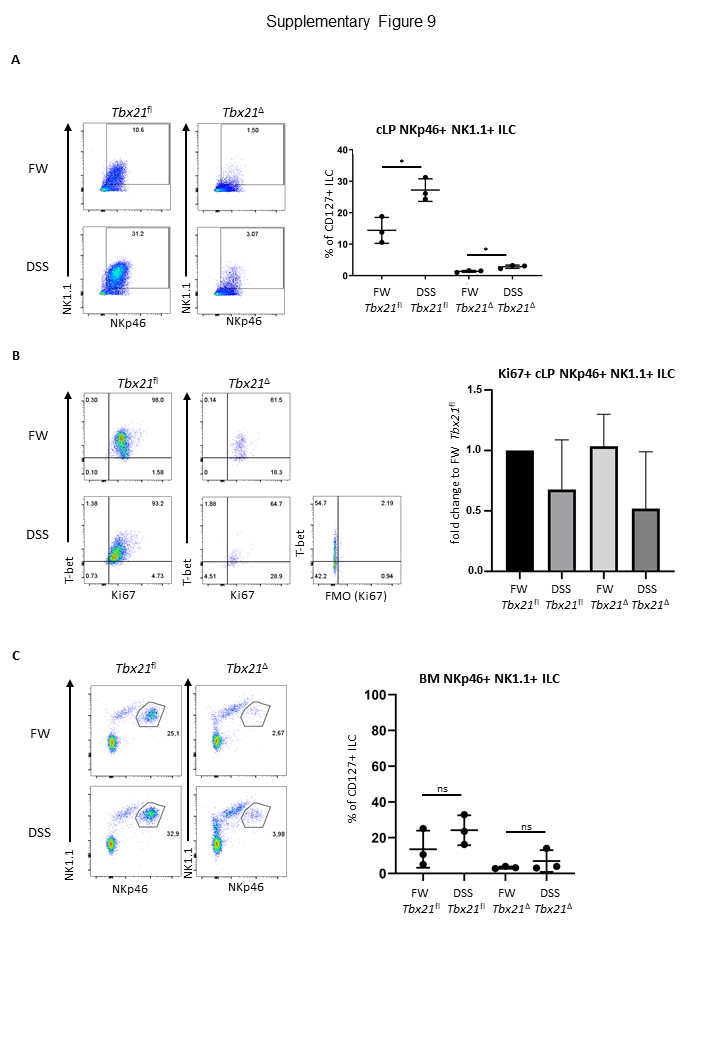
